# Otoferlin requires a free intravesicular C-terminal end for synaptic vesicle docking and fusion

**DOI:** 10.1101/2024.03.20.585901

**Authors:** Didier Dulon, Jacques Boutet de Monvel, Baptiste Plion, Adeline Mallet, Steven Condamine, Yohan Bouleau, Saaid Safieddine

**Author notes:** Authors Contributions: S.S. and D.D. designed the experiments and wrote the paper. J.B. and S.S. performed FRAP, immuno STED microscopy and electron tomography (with A.M). Y.B and B.P. performed confocal immunofluorescence imaging. S.C., Y.B. and D.D. performed ABRs, DPOAEs measurements and hair cell recordings. Declaration of interests: “The authors declare no competing interests.”.

## Abstract

**SUMMARY:** Our understanding of how otoferlin, the major calcium sensor in inner hair cells (IHCs) synaptic transmission, contributes to the overall dynamics of synaptic vesicle (SV) trafficking remains limited. To address this, we generated a knock-in mouse model expressing an otoferlin-GFP protein, where GFP was fused to its C-terminal transmembrane domain. Similar to the wild type protein, the GFP-tagged otoferlin showed normal expression and was associated with IHC SV. Surprisingly, while the heterozygote *Otof^+/GFP^* mice exhibited a normal hearing function, homozygote *Otof^GFP/GFP^* mice were profoundly deaf attributed to severe reduction in SV exocytosis. Fluorescence recovery after photobleaching revealed a markedly increased mobile fraction of the otof-GFP-associated SV in *Otof^GFP/GFP^*IHCs. Correspondingly, 3D-electron tomographic of the ribbon synapses indicated a reduced density of SV attached to the ribbon active zone. Collectively, these results indicate that otoferlin requires a free intravesicular C-terminal end for normal SV docking and fusion.

## INTRODUCTION

The mammalian hearing organ, the cochlea, harbors two types of sensory cell, the inner (IHCs) and outer hair cells (OHCs). While the OHCs act as mechanical amplifiers of the organ’s sound vibrations, the IHCs are the primary sensory cells that transduce these vibrations into electrical nerve impulses conveyed to the brain by auditory afferent fibers. This neurotransmission process is triggered by voltage-dependent activation of Ca_V_1.3 calcium channels located at the IHC synaptic active zones (Brandt et al., 2005; Vincent et al., 2018), leading to the exocytosis of glutamatergic vesicles whose release content activates postsynaptic AMPA receptors (Glowatzki and Fuchs, 2002; Rutherford et al., 2023). The fusion of the synaptic vesicles in IHCs takes place at a highly specialized structure, the synaptic ribbon, an electron-dense structure of submicron diameter that marks the center of the synaptic active zone and to which synaptic vesicles are attached (Smith and Sjostrand, 1961; Lenzi and von Gersdorff, 2001; Lenzi et al., 2002; Safieddine et al., 2012). In contrast to the action potential-initiated bursts of transmitter release occurring at CNS neuronal synapses, IHC ribbon synapses can sustain extremely high rate of transmitter release. This rate is continuously modulated in response to sound-induced membrane depolarization (Goutman and Glowatzki, 2007). Prior to being primed for fusion, IHC synaptic vesicles undergo a sequence of steps mirroring those documented in CNS synapses, including trafficking and docking to the active zone through interactions with SNARE proteins (Safieddine et al., 2012; Calvet et al. 2022). Synaptic vesicle exocytosis, activated by Ca^2+^ entry through nearby Ca^2+^ channels, involves two functionally distinct pools of vesicles, namely a readily releasable pool (RRP) composed of primed vesicles located below the ribbon and directly apposed to the active zones, and a sustained releasable pool (SRP) of vesicles likely attached to the ribbon or in its close vicinity (Moser and Beutner, 2000; Vincent et al., 2017).

Otoferlin, a six C2 domain protein defective in a recessive form of human pre-lingual deafness DFNB9 (Yasunaga et al., 1999), acts as an essential Ca^2+^ sensor for RRP and SRP exocytosis in IHCs (Roux et al., 2006; Michalski et al., 2017, Leclère and Dulon, 2023). Upon binding to calcium, otoferlin interacts with the IHC membrane fusion machinery and the lipid membranes, thereby triggering synaptic vesicle fusion with the pre-synaptic plasma membrane. Interestingly, otoferlin is enriched, but not restricted, in the membrane of IHC synaptic vesicles and in the pre-synaptic plasma membrane (Roux et al., 2006; Uthaiah and Hudspeth, 2010; Vogl et al. 2015), suggesting that the protein could also be involved in synaptic vesicle trafficking to the active zone (Heidrych, 2008), in the RRP replenishment (Pangrsic et al. 2010), in the maintenance of IHC calcium influx (Pangrsic et al., 2010; Vincent et a., 2017), in the priming process (Vogl et al., 2015), or even in endocytosis (Heidrych et al., 2009; Tertrais et al., 2019).

Remarkably, otoferlin is a tail-anchored protein through a single-spanning C-terminal transmembrane domain (TMD), whose role remain unknown. Mutations in exon 48 encoding the otoferlin TMD, leading to hearing impairment, underline the importance of this domain (Vogl et al., 2016; Iwasa et al., 2019; Vona et al., 2020; Santarelli et al., 2021). To shed further light on the role of otoferlin in the IHC synaptic vesicle cycle and the importance of its TMD, we engineered a knock-in mouse model expressing an otoferlin-GFP fusion gene, in which EGFP was fused to the intravesicular carboxy-terminal end of its TMD (Fig.1). We thoroughly examined these mice using a comprehensive approach, including electrophysiology, measurement of inner hair cell (IHC) membrane capacitance, Two-photon fluorescence recovery after photobleaching (FRAP) experiments, confocal and STED microscopy, as well as 3D electron tomographic reconstruction of the ribbon synapses. Surprisingly, we found that, despite the GFP-tagged otoferlin showing normal expression and subcellular distribution, the homozygote *Otof ^GFP/GFP^* mice were profoundly deaf, while the heterozygote *Otof ^+/GFP^* mice had normal hearing. We show that the hearing deficit in the *Otof ^GFP/GFP^*mice is explained by a severe reduction in the rates of docking and fusion of the synaptic vesicles at the ribbon active zone. Our study reveals the essential role of the otoferlin TMD in these processes.

**Figure 1:**
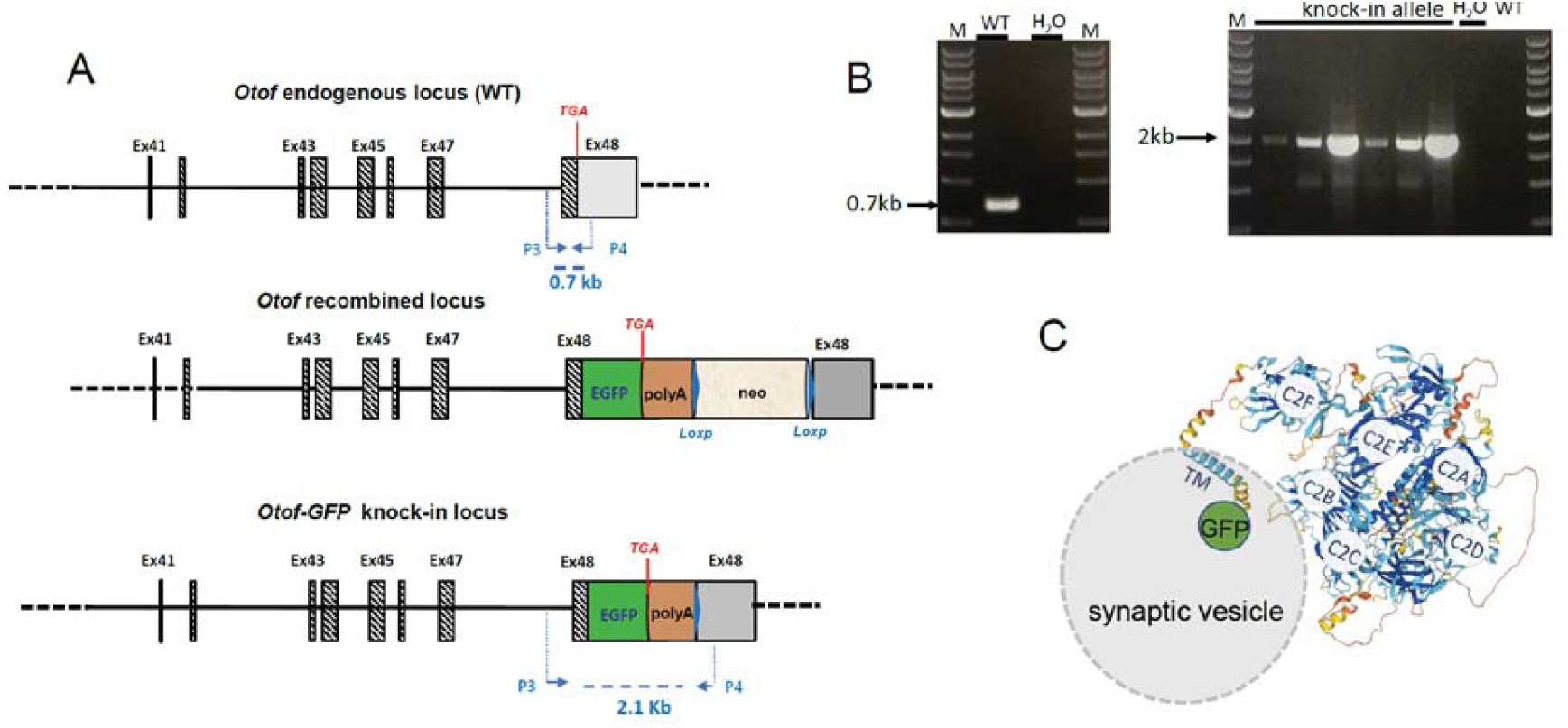
Engineering a mouse line carrying EGFP-tagged *Otof* gene. A) Schematic representation of the *Otof* endogenous wild-type (WT), recombined knock-in allele before and after Cre-recombination, are shown. The last 8 of the 48 exons of Mus musculus otoferlin gene (*Otof*) are noted according to NCBI sequence identifier: NM_001100395.1. B) The primer pair P3 and P4 is tailored to specifically detect the deletion of the wild-type allele (WT). These primers amplify a PCR product of 0.7 Kb for the WT allele and 2.1 Kb for the EGFP knock-in allele. C) Schematic drawing representing a GFP-tagged otoferlin attached to a synaptic vesicle. The C-terminal part of otoferlin after the transmembrane domain (TM), to which is attached GFP, is proposed to be intravesicular. The 3D structure of otoferlin sequence was obtained using Alphafold prediction analysis (Jumper et al., 2021).

## RESULTS

### Generation and characterization of Otoferlin-GFP knock in mouse

In order to elucidate the function of the otoferlin transmembrane domain and investigate the in vivo mobility of synaptic vesicles in live IHCs, we generated a knock-in mouse expressing an otoferlin-GFP fusion protein from the endogenous otoferlin genomic locus using homologous recombination in mouse embryonic stem cells (Otof-GFP KIs). The GFP tag was fused to the C-terminus by placing the enhanced GFP (EGFP) cDNA within the exon 48 just before the stop codon of the otoferlin open reading frame followed by a neomycin resistance cassette in the 3’ untranslated region. Correctly targeted homologous recombinant ES clones were identified by southern blot and used to generate chimeric mice that transmitted the targeted allele to their offspring. RT-PCR experiments confirmed the presence of a recombinant *Otof*-EGFP fused to the 48 exon just before the stop codon (Fig. 1A-B). The otoferlin TMD is followed by 13 amino acids forming a short hydrophilic segment likely to be intravesicular, to which the GFP was attached (Fig. 1C). To eliminate possible deleterious effects of the neomycin resistance cassette, heterozygous otoferlin-GFP mice were first crossed with PGK-Cre mice that expressed the Cre recombinase under control of the phosphoglycerate kinase (PGK) promoter, and then intercrossed to produce homozygous mice. The heterozygote and homozygote mice, referred to below as *Otof ^+/GFP^ and Otof ^GFP/GFP^* mice, respectively, were born at the expected Mendelian ratio, and both were viable and fertile. These mutant mice showed no obvious morphological defect and had overall normal behavior.

### *Otof ^GFP/GFP^* mice are profoundly deaf

Auditory brainstem responses (ABRs), reflecting the activity of the auditory neurons in the central auditory pathway in response to sound stimuli, were recorded in *Otof ^+/+^ (WT)*, *Otof ^+/GFP^* and *Otof ^GFP/GFP^* littermates (Fig. 2). *Otof ^+/GFP^* mice (n = 9) produced characteristic ABR waveforms similar to WT mice (n = 8), with thresholds ranging between 20 and 45 dB SPL, whereas *Otof ^GFP/GFP^*mice (n = 14) showed no detectable ABR, even for stimulus intensities above 90 dB SPL (Fig. 2A,B,C). The similarity between the distortion product otoacoustic emissions (DPOAEs) observed in *Otof ^GFP/GFP^* and WT mice indicated that the profound deafness in *Otof ^GFP/GFP^* mice is likely associated with a defect in the synaptic transmission of IHCs (Fig.2D). We concluded from these results that the homozygote *Otof ^GFP/GFP^* mice have an auditory phenotype similar to that previously described in *Otof KO* mice, characteristic of an IHC synaptopathy (Roux et al., 2006).

**Figure 2:**
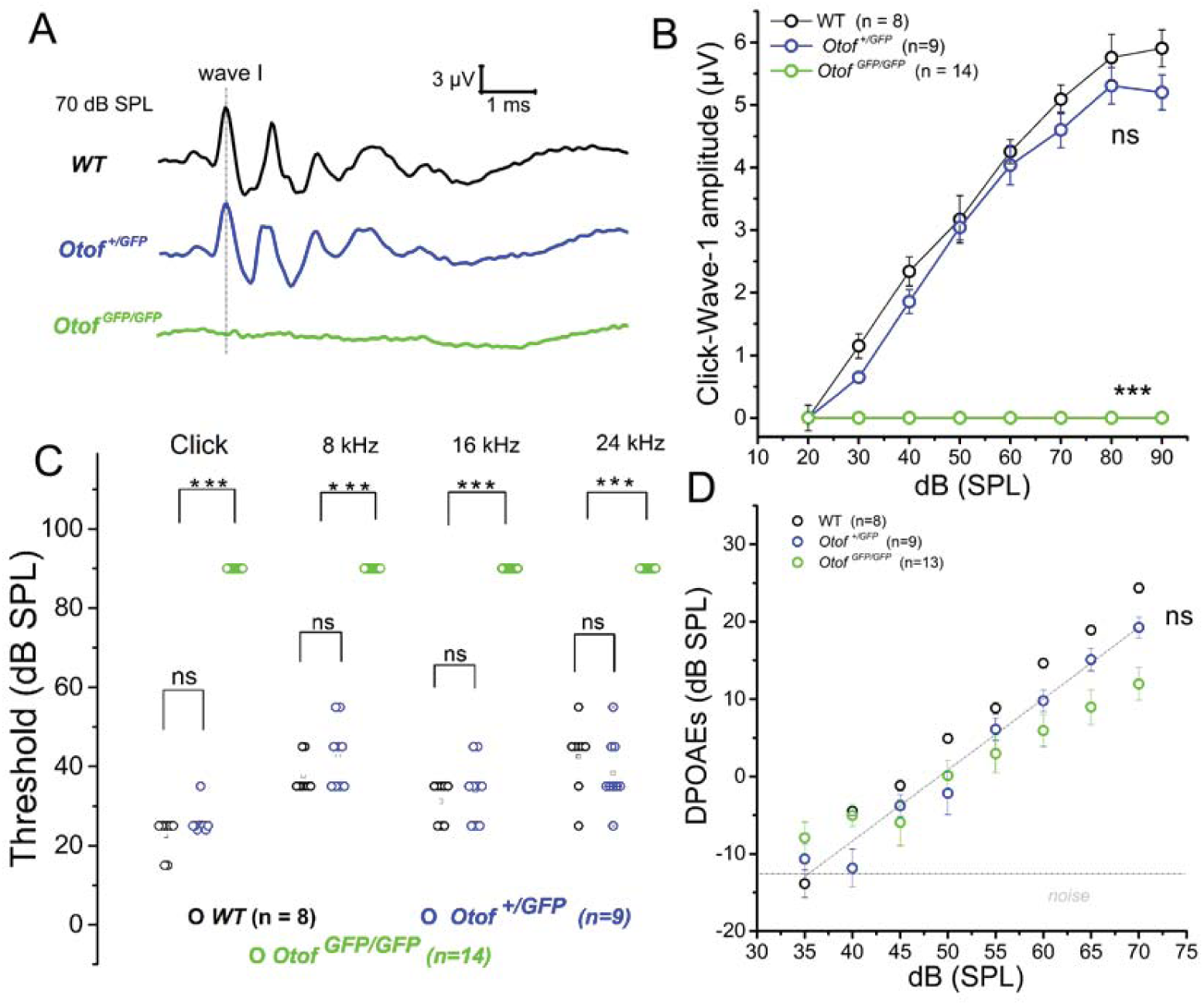
*Otof ^+/GFP^* mice have normal hearing while *Otof ^GFP/GFP^* are profoundly deaf. A) Representative examples of click-ABR waves recorded at 70 dB SPL B) Comparative click-wave I amplitude as a function of sound level. C) Comparative of auditory thresholds of click and tone-evoked ABR. D) Distortion product otoacoustic emissions (DPOAEs; 2f_1_-f_2_ with f_1_ = 12.73 kHz and f_2_ = 15,26 kHz) as a function of sound intensity. ABRs and DPOAEs were recorded in P25-P30 mice. Asterisks indicate significant difference with p<0.05 as compared to WT (*Otof ^+/+^* mice) while ns indicates not significantly different (two-way ANOVA, Tukey’s multiple comparison test).

### Otoferlin-GFP colocalizes with native otoferlin and synaptic vesicles in IHCs

Neither *Otof ^+/GFP^*nor *Otof ^GFP/GFP^*mice exhibited any noticeable morphological abnormality in the organ of Corti, whose sensory epithelium displayed its characteristic organization composed of three rows of OHCs and one row of IHCs of normal appearance. Similar GFP fluorescence levels within the sensory hair cells from both genotypes were also observed when assessed by confocal or two-photon microscopy. This could be observed whether directly *ex vivo* in “live” freshly dissected organs of Corti, maintained in artificial perilymph (Fig. 3A,B), or after chemical fixation (Fig. 4A,B). Otoferlin-GFP signal was detected in IHCs and OHCs from embryonic day 16 (E16) on, and was no longer detectable in the OHCs by the onset of hearing around postnatal day 12 (P12) in mice, and onwards. This pattern aligns well with previous reports on native otoferlin, suggesting that the GFP-tag did not disrupt the developmental regulation of otoferlin expression (Roux et al., 2006; Beurg et al., 2008; Roux et al., 2009). Dual immunolabeling of the heterozygote mouse cochleas for native otoferlin and otoferlin-GFP demonstrated that both proteins exhibited comparable subcellular distribution (Fig. 4B). As expected, the expression of otoferlin-GFP was also observed in vestibular tissues at P2 and P30, with both type I and type II hair cells displaying strong fluorescence (Fig. 3C). These results are consistent with findings from previous vestibular immunolocalization studies (Shug et al., 2006; Dulon et al., 2009).

**Figure 3:**
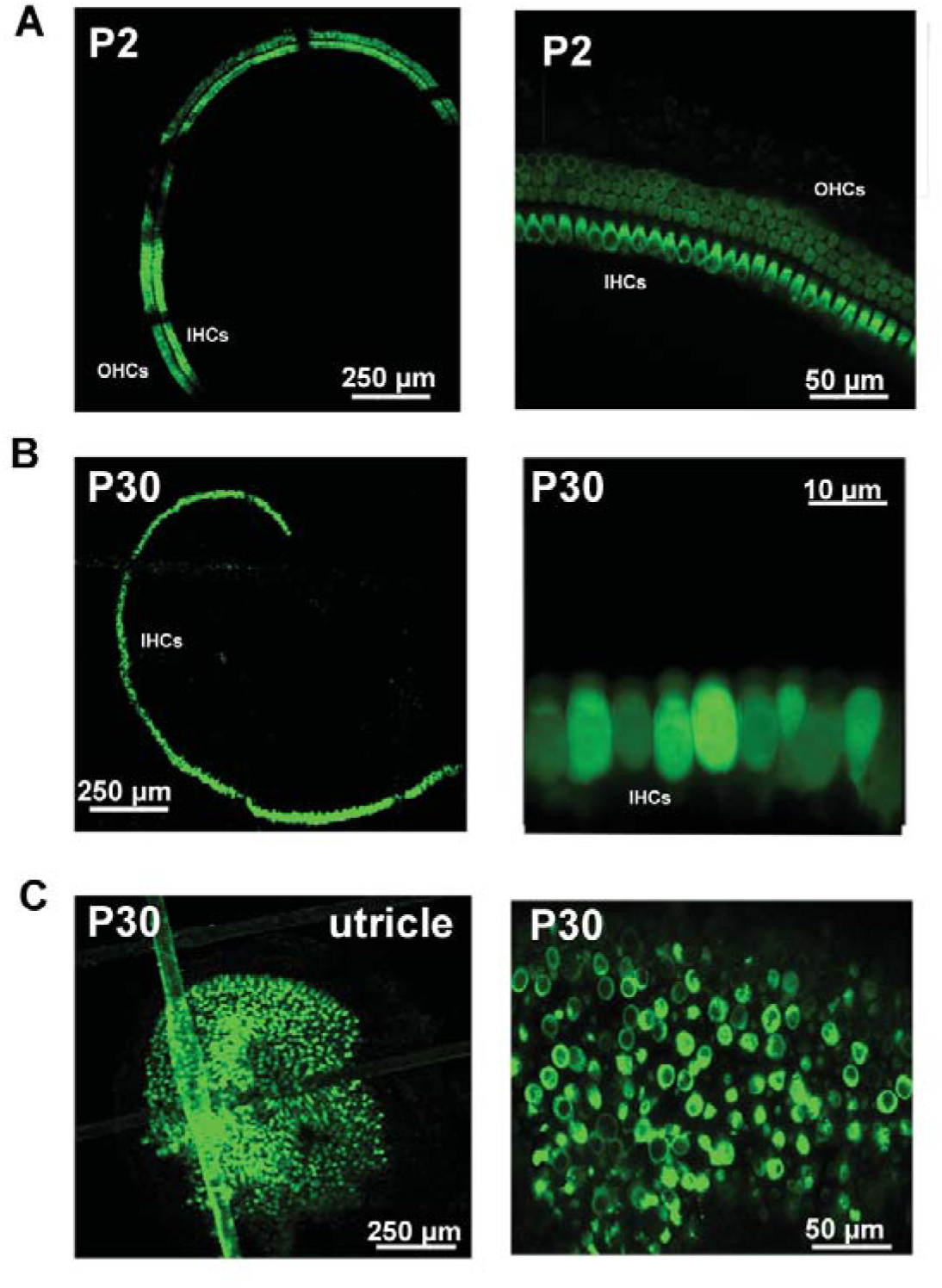
Expression of otoferlin-GFP in freshly dissected living inner ear tissue from Otof ^GFP/GFP^ mice. A) Confocal fluorescence imaging of a living surface preparation from an organ of Corti of a pre-hearing P2 mouse showing a bright GFP fluorescence in both IHCs and OHCs B) Surface preparation of an organ of Corti from a post-hearing mature P30 mouse showing otoferlin-GFP fluorescence restricted to IHCs. C) Living surface preparation of a vestibular utricular organ from a P30 mouse, showing otoferlin-GFP expression in vestibular hair cells.

**Figure 4:**
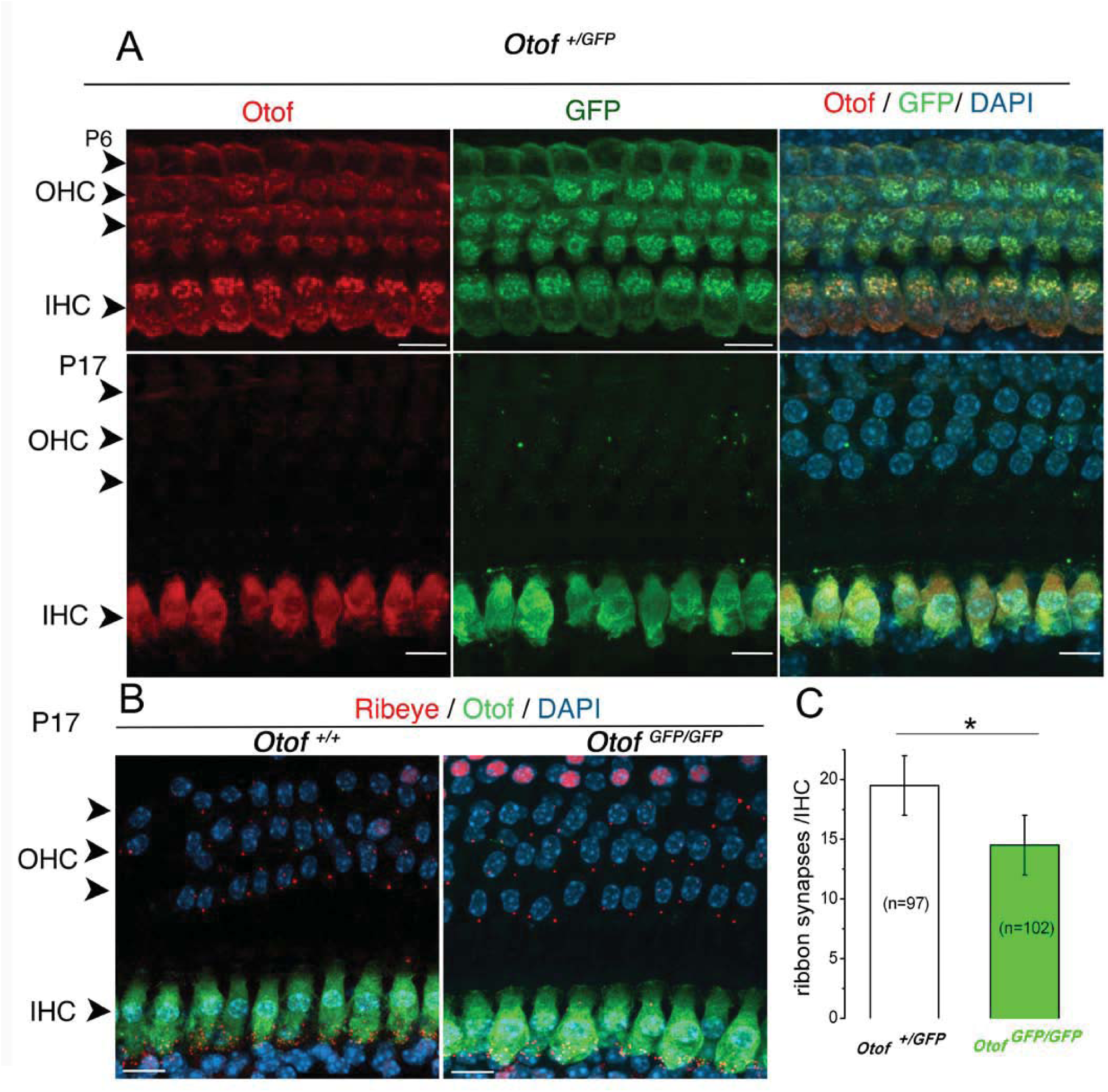
Otoferlin-GFP colocalizes with native otoferlin. A) Confocal immunofluorescence imaging of the native otoferlin (red) and otoferlin-GFP (anti-GFP) in fixed surface preparations of organs of Corti in P6 and P17 *Otof ^+/GFP^* mice. Note the loss of otoferlin-GFP expression, as the native otoferlin, in P17 OHCs. B) Confocal immunofluorescence imaging of otoferlin-GFP and CtBP2 (ribeye, in red) in *Otof^+/+^* mice and *Otof ^GFP/GFP^* mice. C) Histogram showing a significant synaptic ribbon loss in IHCs from P17 *Otof ^GFP/GFP^* mice (p<0.05; unpaired-t-test).

In order to verify that the GFP-tag did not disrupt the subcellular distribution of otoferlin, we examined the co-localization of otoferlin-GFP in the IHCs with the VGLUT3 specific vesicular glutamate transporter type 3 (VGLUT3), which localizes specifically to the IHC synaptic vesicles (Ruel et al., 2008; Seal et al., 2008; Obholzer et al., 2008). The patterns of otoferlin-GFP fluorescence and anti-VGLUT3 immunofluorescence were comparable under conventional confocal microscopy (Fig. 5). The precise subcellular distribution of otoferlin-GFP and its colocalization with VGLUT3 in the IHCs were further examined by stimulated emission depletion (STED) microscopy, which provides a resolution surpassing the optical diffraction limit (Hell and Wichmann 1994) and has been successfully used to visualize individual synaptic vesicles in hypocampal neurons (Willig et al, 2006; Westphal et al, 2008). When visualized in the intracellular IHC regions by dual-color STED microscopy (Lauterbach et al, 2013), the distribution of both DY485XL-stained VGLUT3 proteins and Atto565-stained GFP formed numerous fluorescent spots of a diameter ranging 80-100 nm. These spots could be readily distinguished within a single optical plane section, thus likely localizing individual synaptic vesicles. Examination of the nearest-neigbor distances between GFP-Atto565 and VGLUT3-DY485XL spots validated the robust colocalization between the two populations of spots (Fig. 5).

**Figure 5:**
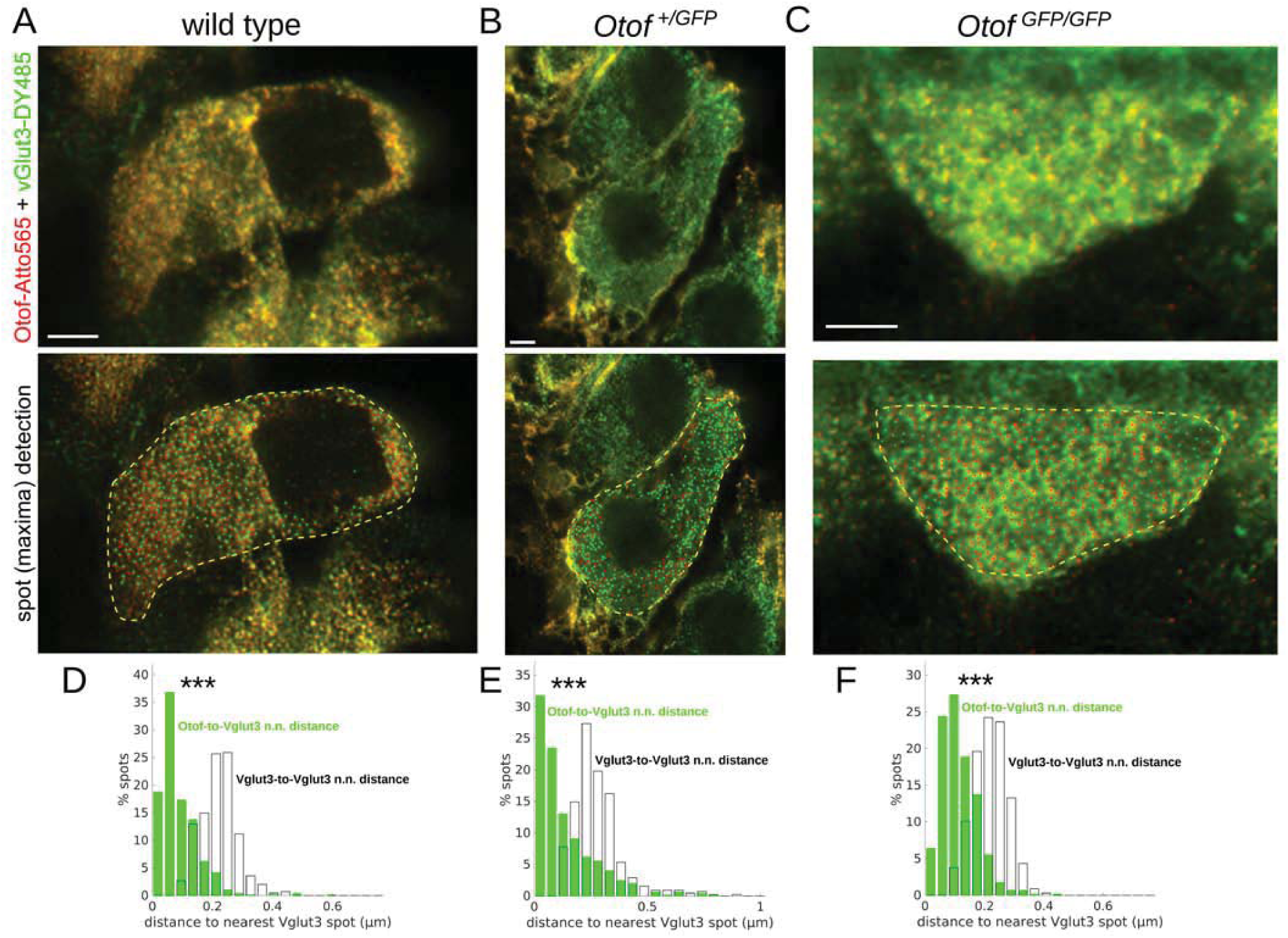
Colocalization of otoferlin-GFP and VGLUT3. (A-C) Upper panels: Representative STED images of individual P15 IHCs in whole-mount preparations from wild type (A), *Otof ^+/GFP^* (B) and *Otof ^GFP/GFP^* (C) cochleas immunolabeled for otoferlin and VGLUT3 (A), or for otoferlin-GFP and VGLUT3 (B and C). Scalebars, 2µm. Lower panels: The same images after wavelet denoising and deconvolution, with the positions of the resolution-limited Atto565 and DY485 spots displayed in red and green, respectively, within the delineated IHC (yellow dashed contour). (D-F) Plots of the distributions of nearest neighbor distances between otoferlin spots and their corresponding closest VGLUT3 spots (green-filled histogram), or between VGLUT3 spots and their corresponding closest VGLUT3 spots (black histogram without fill), for the wild-type (D), the *Otof +/GFP* (E), and the *Otof GFP/GFP* (F) IHCs. In all cases, the otoferlin-to-VGLUT3 nearest neighbor distance distribution display a marked leftward shift (toward smaller distances) compared to the VGLUT3-to-VGLUT3 nearest neighbor distance distribution (p<10-4, Kolmogorov-Smirnov test), supporting the colocalization of otoferlin-Atto565 with VGLUT3-DY485 spots in these IHCs.

Altogether, these observations demonstrate that the GFP-tag fused to the otoferlin TMD on its intravesicular side does not interfere with the spatiotemporal expression pattern and subcellular localization otoferlin. Otoferlin-GFP appears properly targeted to the synaptic vesicles of IHCs, and its distribution within the cochlea and in vestibular hair cells was indistinguishable from that of the endogenous otoferlin (Roux et al. 2006, 2009; Dulon et al., 2009).

### IHCs of *Otof ^+/GFP^* mice exhibited normal synaptic exocytosis

To verify whether IHC synaptic vesicle exocytosis was affected by the presence of otoferlin-GFP, we performed *ex-vivo* whole-cell patch clamp recordings of membrane capacitance in IHCs from post-hearing P18-P20 heterozygote *Otof ^+/GFP^* mice. At holding membrane potential of −70 mV, in absence of stimulation, the cell membrane capacitance (a proxy of resting cell size) of heterozygote *Otof ^+/GFP^* IHCs (10.97 ± 0.26 pF; n = 22) was found similar to WT IHCs (10.61 ± 0.16 pF; n = 22; SEM, unpaired t-test, p = 0.26; Fig. 6A), indicating a normal resting balance between exocytosis and endocytosis. During voltage-step depolarization, *Otof ^+/GFP^* IHCs displayed Ca^2+^ current with amplitude and voltage-dependence similar to WT IHCs (Fig. 6B). We then characterized the kinetics of Ca^2+^-evoked exocytosis (ϕλC_m_) of the RRP by depolarizing IHCs from −80 mV to −10 mV with increasing duration from 10 to 80 ms (Fig. 6C). The plot of ϕλC_m_ as a function of time indicated an exponential rise with similar time constants in WT (1 = 26 ± 8 ms; n = 12) and in heterozygote *Otof ^+/GFP^* IHCs (29 ± 4 ms; n = 12; SEM, unpaired t-test, p = 0.78). The ϕλC_m_ responses reached a similar plateau within 80 ms at 22.7 ± 2.3 fF and 23.0 ± 2.0 fF in WT and *Otof ^+/GFP^*, respectively (Fig. 6C). Overall, these results are consistent with the normal ABRs found in heterozygote *Otof ^+/GFP^* mice, and confirm that a single allele expression of otoferlin-GPF, together with an expression of native otoferlin by the other WT allele, do not modify synaptic vesicular release in IHCs.

**Figure 6:**
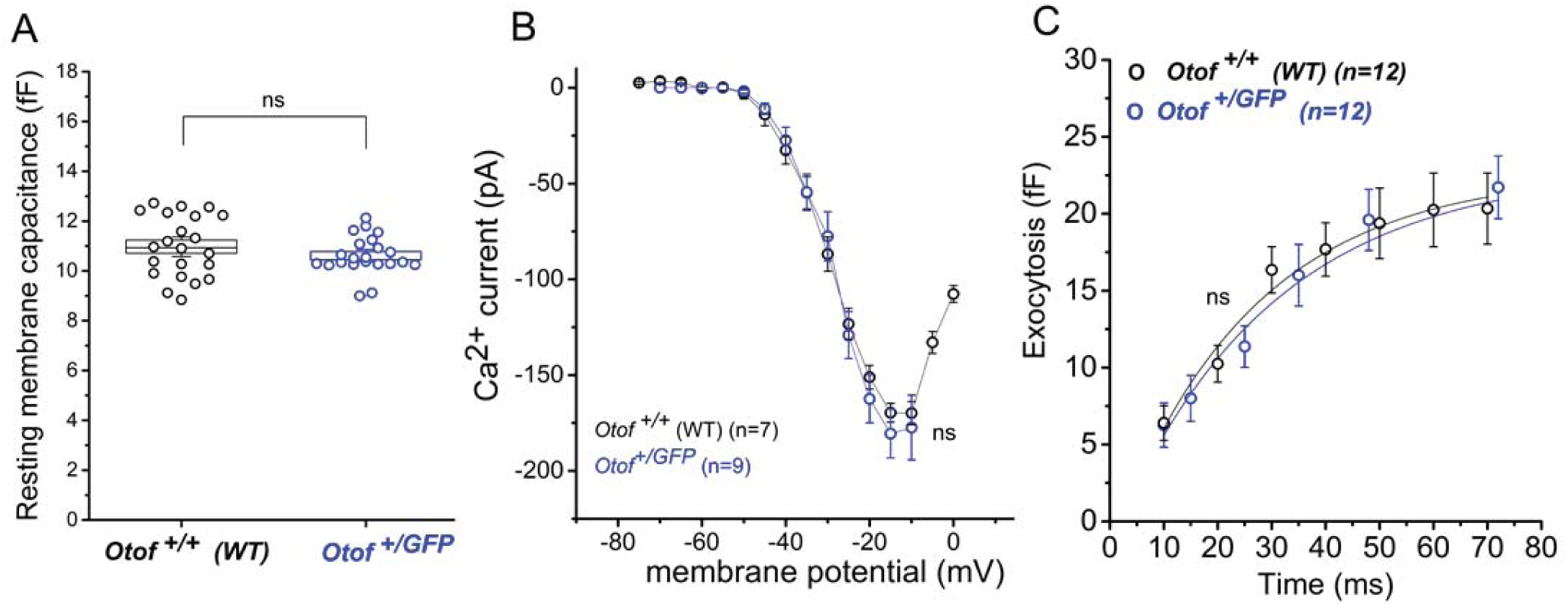
Ca^2+^ currents and exocytosis are normal in IHCs of P14-P18 *Otof ^+/GFP^* mice. A) Resting membrane capacitance of IHCs, measured at V_h_ = −70 mV, is comparable in WT and *Otof ^+/GFP^* mice (p=0.25; unpaired-t-test). B) Ca^2+^ currents evoked during 25 ms voltage-steps at various potentials and (C) kinetics of exocytosis (constant voltage-step from −80 mV to −10 mV with increasing time duration) in *Otof ^+/GFP^* IHCs were comparable to WT. Values are mean with SEM and ns indicates not significantly different (two-way ANOVA, Tukey’s multiple comparison test).

### Synaptic exocytosis in *Otof ^GFP/GFP^* IHC is severely impaired

The profound deafness with normal DPOAEs that characterize the *Otof ^GFP/GFP^* mice strongly indicates a malfunction in IHC synaptic transmission. We therefore investigated the IHC synaptic exocytotic function in these mice before and after the onset of hearing. The voltage-gated Ca^2+^ influx and exocytotic membrane capacitance changes were monitored in pre-hearing (P8) and post-hearing (P18-P20) from both *Otof ^+/GFP^* and *Otof ^GFP/GFP^* IHCs. At a holding potential of −70 mV, in absence of stimulation, the cell resting membrane capacitance was found significantly smaller in *Otof ^GFP/GFP^* IHCs at both developmental stages (8.71 ± 0.21 pF at P8, n=8, and 8.84 ± 0.22 at P14-P18, n = 15, SEM) compared to *Otof ^+/GFP^* IHCs (9.77 ± 0.22 pF at P8, n = 8, and 10.61 ± 0.16 pF at P14-P18, n = 22; SEM, unpaired t-test, p < 0.05 for both comparisons). This difference in resting membrane capacitance suggested a balance defect between vesicle exocytosis and endocytosis in homozygotes *Otof ^GFP/GFP^* IHCs.

At P8, while Ca^2+^ currents recorded in *Otof ^GFP/GFP^* and in *Otof ^+/GFP^* IHCs were similar in terms of amplitude and voltage-dependence (Fig. S1A), the kinetics and amplitude of the RRP component of exocytosis in P8 *Otof ^GFP/GFP^* IHCs were largely reduced as compared to P8 *Otof ^+/GFP^* IHCs (Fig. S1B). For a brief 20 ms depolarization, that mostly involves vesicles from the RRP, the amplitude of the exocytotic responses averaged 7.8 ± 1.1 fF (n = 7, SEM) in P8 *Otof ^GFP/GFP^* IHCs as compared to 15.5 ± 2.7 fF (n = 7, SEM) in P8 WT IHCs (unpaired t-test; p < 0.05). The SRP of exocytosis, tested by repetitive 100 ms depolarization steps, was also largely affected in the homozygote *Otof ^GFP/GFP^* mutant mouse (Fig. S1C).

A severe reduction in the kinetics of the RRP and SRP exocytosis was also observed in post-hearing P18-P20 *Otof ^GFP/GFP^* IHCs as compared to *Otof ^+/GFP^* IHCs (Fig. 7). Notably, in these P18-P20 homozygote IHCs, peak Ca^2+^ currents (I_Ca_) were reduced by a factor ∼2.5 as compared to *Otof ^+/GFP^* IHCs (52 ± 10 pA (n = 18, SEM) and 133 ± 7 pA (n = 12, SEM), respectively, unpaired t-test, p< 0.05; Fig. 7A). This reduction in the amplitude of the Ca^2+^ currents could explain in part the defect in exocytosis in P18-P20 homozygote *Otof ^GFP/GFP^* IHCs. To probe the fusion process in *Otof ^GFP/GFP^* IHCs independently of the Ca^2+^ channels, we performed intracellular Ca^2+^ uncaging experiments on IHCs loaded with DM-nitrophen using UV-flash photolysis. While Ca^2+^ uncaging produced a similar fast rise in membrane capacitance in *Otof ^+/GFP^* and WT IHCs, this process was markedly reduced in P18 homozygote *Otof ^GFP/GFP^* IHCs (Fig. 7D), indicating that the Ca^2+^-dependent vesicular fusion process was impaired independently of a possible defect in the expression/organization of the Ca^2+^ channels. Overall, these results demonstrate that although otoferlin-GFP, is normally targeted to the synaptic vesicles (Fig. 4 and Fig. 5), it did not function effectively as a Ca^2+^ sensor for exocytosis in the *Otof ^GFP/GFP^* mutants.

**Figure 7:**
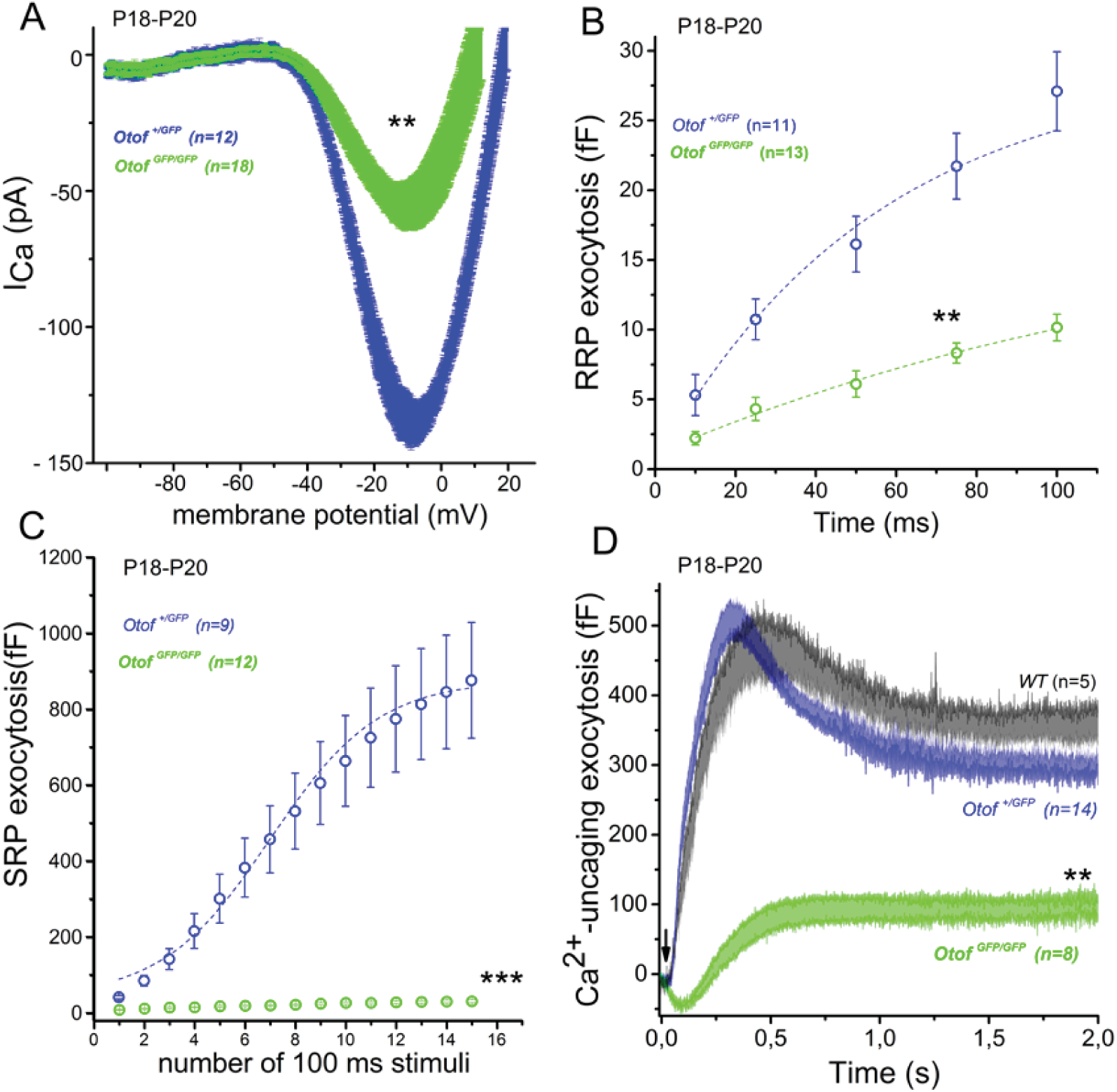
RRP and SRP exocytosis defect in IHCs from P18-P20 *Otof ^GFP/GFP^* mice. A) Mean Ca^2+^ currents recorded in *Otof ^+/GFP^* (blue) and *Otof ^GFP/GFP^* (green) IHCs during a depolarizing voltage-ramp protocol (1 mV/ms) from –90 to +30 mV. Note the reduction in the amplitude of the Ca^2+^ currents in *Otof **^GFP/GFP^***IHCs as compared to *Otof **^+/GFP^*** IHCs (maximum amplitude at −10 mV: 52 ± 10 pA (n = 18) and 133 ± 7 pA (n = 12), respectively, unpaired t-test, p< 0.01). B) Kinetics of RRP exocytosis evoked during brief voltage steps from –80 to –10 mV, with durations increasing from 10 to 100 milliseconds, were largely reduced in IHCs from *Otof* ***^GFP/GFP^*** IHCs (two-way ANOVA, p<0.01). C) SRP exocytosis evoked during long-lasting voltage steps from –80 to –10 mV, with durations increasing from 100 to 1500 ms (100 ms increment), was completely abolished in *Otof* ***^GFP/GFP^*** IHCs. D) Mean exocytotic response curves evoked by intracellular Ca^2+^ uncaging. Note the strong reduction in the rate and amplitude of *Otof **^GFP/GFP^*** IHCs responses as compared to WT (black) and *Otof ^+/GFP^* (blue) IHCs. Values are mean with SEM and asterisks indicate significant difference with p<0.05 (two-way ANOVA).

### The IHCs of *Otof ^GFP/GFP^*mice display an abnormally high mobile fraction of synaptic vesicles

To compare the synaptic vesicle dynamics at the IHC synaptic zone of the normally hearing P18-P20 *Otof ^+/GFP^* mice and the deaf P18-P20 *Otof ^GFP/GFP^ mice*, otoferlin-GFP associated vesicles were visualized using two-photon microscopy at the basolateral synaptic region below the cell nucleus, and their mobility was assessed by FRAP.

In *Otof ^+/GFP^* IHCs, the fraction of post-bleach recovery averaged 23.2 ± 2.0 % with a coefficient of diffusion of *D* = 0.33 ± 0.05 μm^2^/s (n = 62, SD, Fig. 8A,8D). These results suggested that a significant, though relatively modest fraction of the otoferlin-GFP proteins were mobile. FRAP recovery curves obtained in similar conditions in the IHCs from *Otof ^GFP/GFP^*mice revealed a 60% increase in the mobile fraction of synaptic vesicles (37.5 ± 2.8 %, SD, n = 55, p < 10^-4^, t-test) as compared to the *Otof ^+/GFP^* mice (Fig. 8B,8D). This large mobile fraction in the *Otof ^GFP/GFP^* IHCs showed however a similar coefficient of diffusion (*D* = 0.27 ± 0.04 μm^2^/s; SD) as observed in *Otof ^+/GFP^* IHCs (p =0.36, t-test). These results suggested that a large pool of synaptic vesicles had weaker interactions with docking sites in *Otof ^GFP/GFP^* IHCs as compared to *Otof ^+/GFP^* IHCs. The coefficient of diffusion of this larger mobile pool was however unaffected.

**Figure 8:**
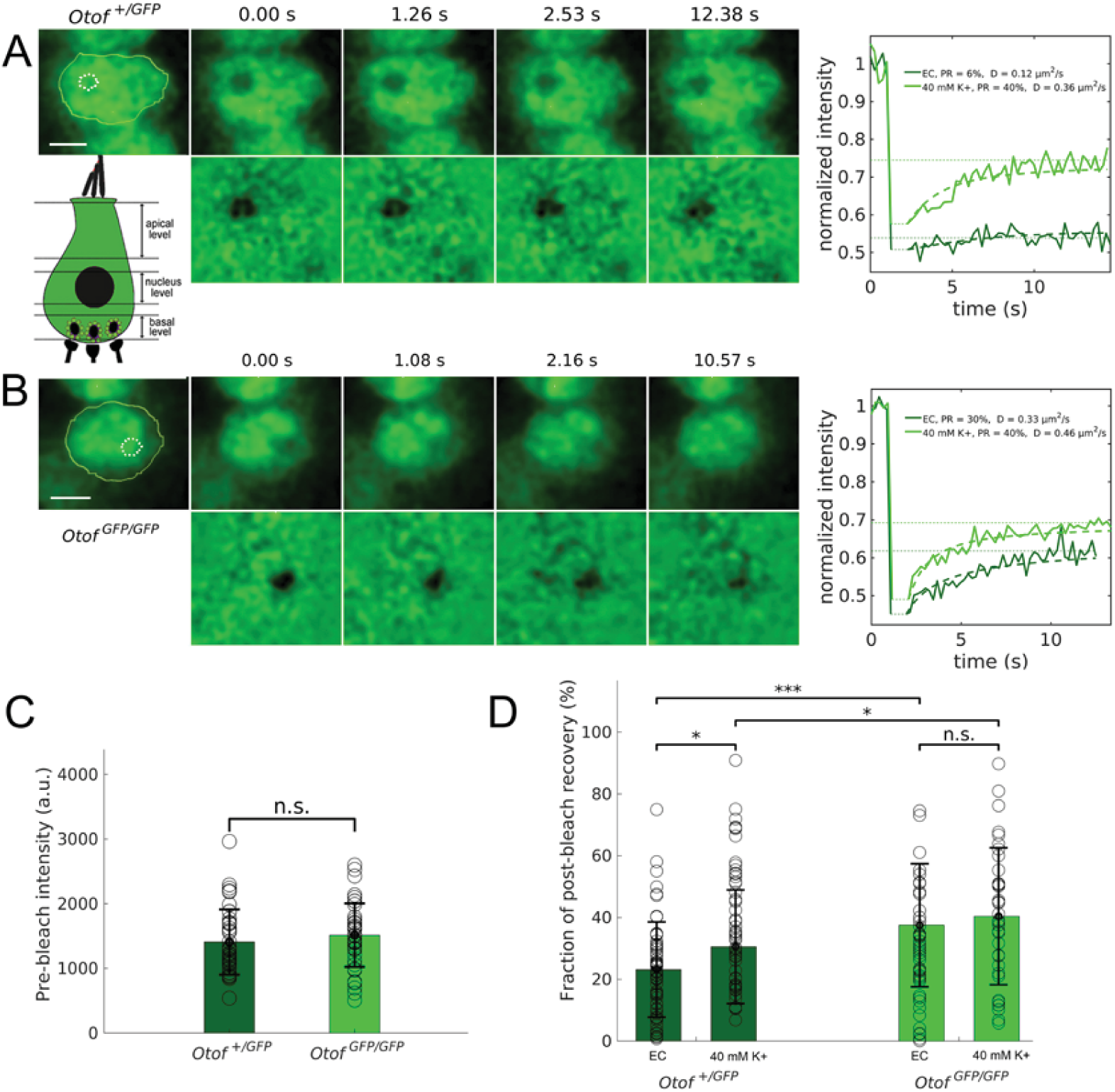
FRAP experiments show a larger mobile fraction in *Otof ^GFP/GFP^* IHCs. Examples of two-photon FRAP experiment on an *Otof ^+/GFP^* IHC (A) and an *Otof ^GFP/GFP^* IHC (B), following a bleach applied at the basal synaptic region of the cell. Shown on the left of each panel are a series of two-photon images acquired prior to the bleach, just after the bleach (t = 0s), and at various times during the fluorescence recovery phase. The bleached spot and the basal cell body are delineate on the pre-bleach images (white and yellow dashed contours, respectively). The lower-left inset in (A) indicates the approximate location of the ∼1μm thick optical section where the bleach was applied. Plotted on the right are normalized fluorescence recovery curves, showing the mean intensity measured within the bleached spot as a function of time, under control conditions in extracellular medium (EC, dark green curves), or while the cells were being depolarized by application of 40 mM K+ to the medium (light green curves). Solid curves represent experimental data, while the dashed curves are best fits to a simulation of the 2D-diffusion equation in the plane of the image (performed as in (Boutet de Monvel et al 2006)). (C) Quantification of the intensity measured within the bleached spot prior to the applied bleach, showing that the GFP signal did not differ significantly in *Otof ^+/GFP^* and in *Otof ^GFP/GFP^* IHCs. (D) Quantification of the recovery fraction (mobile fraction of synaptic vesicles) in *Otof ^+/GFP^*and *Otof ^GFP/GFP^* IHCs, as measured under control conditions (EC), or during 40 mM K+ stimulation. Note the pronounced increase in the mobile fraction measured in the homozygote mutant under control conditions (EC). The mobile fraction of IHC synaptic vesicles was also observed to increase during the K+ stimulation, although this increase was significant only in *Otof ^+/GFP^* IHCs, and remained marginal in *Otof ^GFP/GFP^* IHCs.

To test whether stimulation affected the mobility of synaptic vesicles, we depolarized the IHCs by applying an extracellular solution of 40 mM K^+^ and 2.5 mM Ca^2+^. The mobile fraction of otoferlin-GFP increased to 30.6 ± 2.8% (n=44, SD, p=0.03, t-test) in presence of 40 mM K^+^ in *Otof ^+/GFP^* (Fig. 8A,8D). This increased fraction of mobile vesicle suggested that rising intracellular Ca^2+^ decreased otoferlin-GFP intracellular interactions with docking sites. Using a similar K^+^ depolarization protocol in homozygote *Otof ^GFP/GFP^* IHCs, we found that the mean fraction of mobile otoferlin-GFP also slightly increased, but not significantly to 40.4 ± 3.8 % (n = 25; SD, p = 0.53). These results suggested that intracellular Ca^2+^-increase had a positive effect on the mobile fraction of otoferlin-GFP associated synaptic vesicles in the heterozygote mutants, while it had a marginal effect in the homozygotes.

### Reduced number of ribbon synapses in IHC of *Otof ^GFP/GFP^*

The IHC ribbon synapses of P18 *Otof ^GFP/GFP^* and *Otof ^+/GFP^* mice were examined by triple immunofluorescence confocal microscopy, using an antibody directed against the protein CtBP2 (identical to the Ribeye B domain, a major structural components of the ribbon; Schmitz et al. 2000), an antibody against glutamate receptor subunits 2 (GluR2), which are associated with the dendrites of afferent neurons, and an antibody against the Ca_v_1.3 channels that govern IHC vesicle fusion. Normal ribbon synapses were defined by the presence of closely juxtaposed Ribeye, GluR2 and Cav1.3 fluorescent spots (Sergeyenko et al., 2013; Roux et al., 2006). We quantified the number of ribbon synapses (including the ribbon and an associated calcium channel-populated region) using 3D reconstructions and found a mean of 19.5 ± 2.5 ribbons synapses (n = 97; SD) per IHC in the heterozygote *Otof ^+/GFP^* mice. This number agrees well with previous synaptic ribbon synapse quantification in WT IHCs (Vincent et al. 2014). Similar measurements resulted in significantly lower numbers in the homozygote *Otof ^GFP/GFP^* mice, averaging to 14.5 ± 2.5 (n = 102; SD) ribbon synapses per IHC (Fig. 4C; p<0.05). A similar loss of ribbon synapses was also described in otoferlin KO mice (Tertrais et al., 2019, Roux et al., 2006), suggesting that a normal exocytotic synaptic activity is required for the maintenance of the IHC ribbon synapses.

### The IHC ribbon synapses of *Otof ^GFP/GFP^* display a reduced density of docked vesicles

To establish an anatomical correlate for the exocytosis defect observed in homozygote *Otof ^GFP/GFP^* mice, we studied the ultrastructure of the IHC ribbon synapse and quantified its associated pools of vesicles by 3D electron tomography as previously described (Michalski et al., 2017). For each genotype, nine IHC ribbon synapses from cochlear apical turns of P20 mice were reconstructed. To ensure the accuracy of our quantification of the vesicular pool, only data sets that included the center or the entire ribbon were included from tomographic reconstruction. The mean size (volume) of the synaptic ribbons in *Otof ^+/GFP^* IHCs (3.84 10^-3^ ± 1.71 10^-3^ μm^3^; n=9, SD) and in *Otof ^GFP/GFP^* IHCs (6.43 10^-3^ ± 4.26 10^-3^ μm^3;^ n=9, SD) were not significantly different (unpaired t-test p = 0.12), though the distribution of *Otof ^GFP/GFP^* IHC ribbon size showed a larger spread. For each ribbon, synaptic vesicles were classified into three different pools, according to their position relative to the presynaptic plasma membrane and the ribbon: (i) ribbon-associated vesicles with centers lying within 40 nm of the presynaptic plasma membrane were classified as the presumptive readily releasable pool (RRP; in red); (ii) vesicles lying within 80 nm of the ribbon but not apposed to the presynaptic plasma membrane were classified as the ribbon-attached pool (RAP or presumptive SRP; in green), and (iii) vesicles located between 80 nm and 350 nm from the ribbon surface and not apposed to the presynaptic plasma membrane comprised the outlying pool (OP1 80-100 nm and OP2 100-350 nm; yellow and orange) (Fig. 9A). Since the volume of the ribbons showed substantial variation with a wider distribution in the homozygote *Otof ^GFP/GFP^* IHCs, we plotted the number of ribbon-associated vesicles in the RRP, SRP, and RRP+SRP as a function of the ribbon volume (Fig. 9B,C, and D, respectively). While a strong linear correlation between the ribbon volume and the vesicle number was observed in both genotypes for the RRP and SRP, homozygote *Otof ^GFP/GFP^*IHCs displayed a significantly lower synaptic vesicle density attached to the plasma membrane active zone or to the ribbon (RRP slope: 1946 ± 263 vesicles/μm^3^; SRP slope: 6783 ± 433 vesicles/μm^3^; RRP+SRP slope: 8729 ± 672 vesicles/μm^3^) as compared to the heterozygote *Otof ^+/GFP^* IHCs (RRP slope: 3580 ± 663 vesicles/μm^3^; F_test_, p = 0.0027; SRP slope: 8740 ± 805 vesicles/ μm^3^; F_test_, p = 0.041; RRP+SRP slope: 12320 ± 1316 vesicles/ μm^3^, F_test_, p = 0.022). However, the density of the outlying pool of vesicles around the ribbon was found not significantly different between the heterozygote (OP1: 249 ± 80 vesicles/μm^3^; SD, n=9; OP2: 215 ± 65 vesicles/ μm^3^; SD, n = 9) and homozygote IHCs (OP1: 235 ± 59 vesicles/μm^3^; n=9; SD, unpaired t-test, p = 0.51); OP2: 216 ± 58 vesicles/μm^3^; n = 9; SD, unpaired t-test, p = 0.33; Fig 9E).

**Figure 9:**
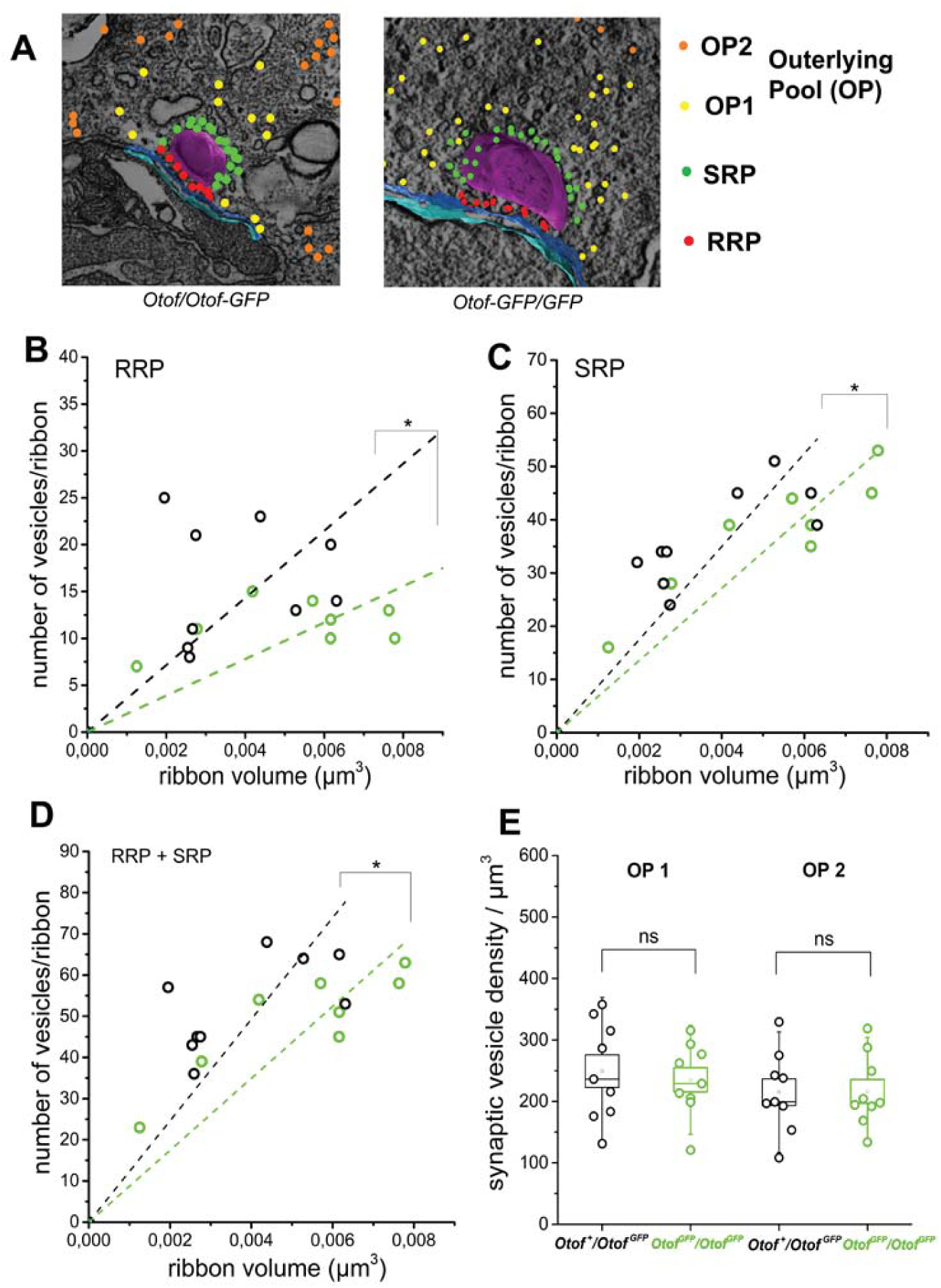
Comparative ultrastructural analysis of the different vesicle pools in *Otof* ^+/GFP^ and *Otof* ^GFP/GFP^ IHC ribbon synapses. A) Representative 3D tomograms of the different pools of synaptic vesicles: presumptive readily-releasable pool (RRP) in red; the ribbon-associated pool or secondary releasable pool (SRP) in green; the outlying pools OP1 (80-100 nm) and OP2 (100-350 nm) are overlaid in yellow and orange, respectively. B-C-D) Comparative ribbon volume relationship with the number of vesicles associated with the RRP and SRP. Note the significant decrease in the number of vesicles for both components in *Otof* ^GFP/GFP^ IHCs (F-test, p < 0.05). E) Mean vesicle densities in the OP1 and OP2 around the ribbon were not significantly different (p = 0.51 and 0.33 respectively, unpaired-t-test).

Overall, our ultrastructural analysis suggested that the complete replacement of the untagged WT form of otoferlin by otoferlin-GFP reduced by nearly half the number of docked vesicle below the synaptic ribbon, while the number of vesicles attached to the ribbon were also reduced by 22%. These results are consistent with the FRAP experiments, suggesting a reduced interaction of the otoferlin-GFP-tagged vesicles with their specific targets at the synaptic ribbon, in the *Otof ^GFP/GFP^* IHCs.

### Intravesicular molecular crowding may explain the deficit in *Otof ^GFP/GFP^* IHCs

It has been suggested that multimerization of synaptotagmin 1, the major a Ca^2+^ sensor for exocytosis at the CNS synapses, is critical for its proper function (Courtney et al., 2021). Likewise, it has been demonstrated that the dimerization of dysferlin, a protein containing six C2 domains and belonging to the ferlin family, through it TMD is crucial for its role as Ca2+ sensor in muscle membrane repair (Xu et al., 2011). To determine whether similarly otoferlin can possibly self-oligomerize through its TMD, we performed structure analysis and representation of the otoferlin TMD using Protter, a web-based tool for interactive protein visualization (Omasits et al., 2014), AlphaFold protein structure data base (Developed by Deep Mind and EMBL-EBI; Jumper et al., 2021; Varadi et al., 2022), and PREDDIMER, a prediction tool for transmembrane a-helical dimer conformations (Polyansky et al., 2012, 2013). Otoferlin was predicted as a single-spanning membrane protein that consists of a large cytoplasmic domain (containing six-C2 domains and the Ca^2+^ binding sites), a juxtamembrane (JM) region (8 amino acids), a transmembrane domain (TMD of 21 amino acids), and a short intravesicular region of 13 amino acids (Fig. 10). Protter analysis predicted a 21 amino acid protein TMD structure containing a leucine zipper-like heptad repeat of amino acids (Fig. 10A-B). This leucin repeat structure has long been recognized to be essential for TMD protein dimerization in several other single-spanning (bitopic) membrane proteins (Landshulz et al., 1988; Gurezka et al., 1999; Cymer and Schneider, 2010; Li et al., 2014). PREDDIMER analysis predicted the possible formation of transmembrane a-helical TMD dimers with excellent probability values (with best *F_SCOR_* value of 3.7; Supplementary table 1). These F*_SCOR_* values are similar to the one generated in several well characterized homodimer proteins whose structures were obtained previously by nuclear magnetic resonance (Polyansky et al., 2013). Together, the above in-silico analyses suggest that the otoferlin TMD works as a dimerization module, which could provide a plausible explanation for synaptic deficit observed in *Otof ^GFP/GFP^* IHCs. Specifically, we propose that tagging the short intravesicular C-terminal tail of TMD with GFP may generate intravesicular molecular crowding and interfer with the assembly-disassembly dynamics of otoferlin TMD dimers, thereby affecting vesicle docking and fusion at the IHC active zone (Fig. 10C).

**Figure 10:**
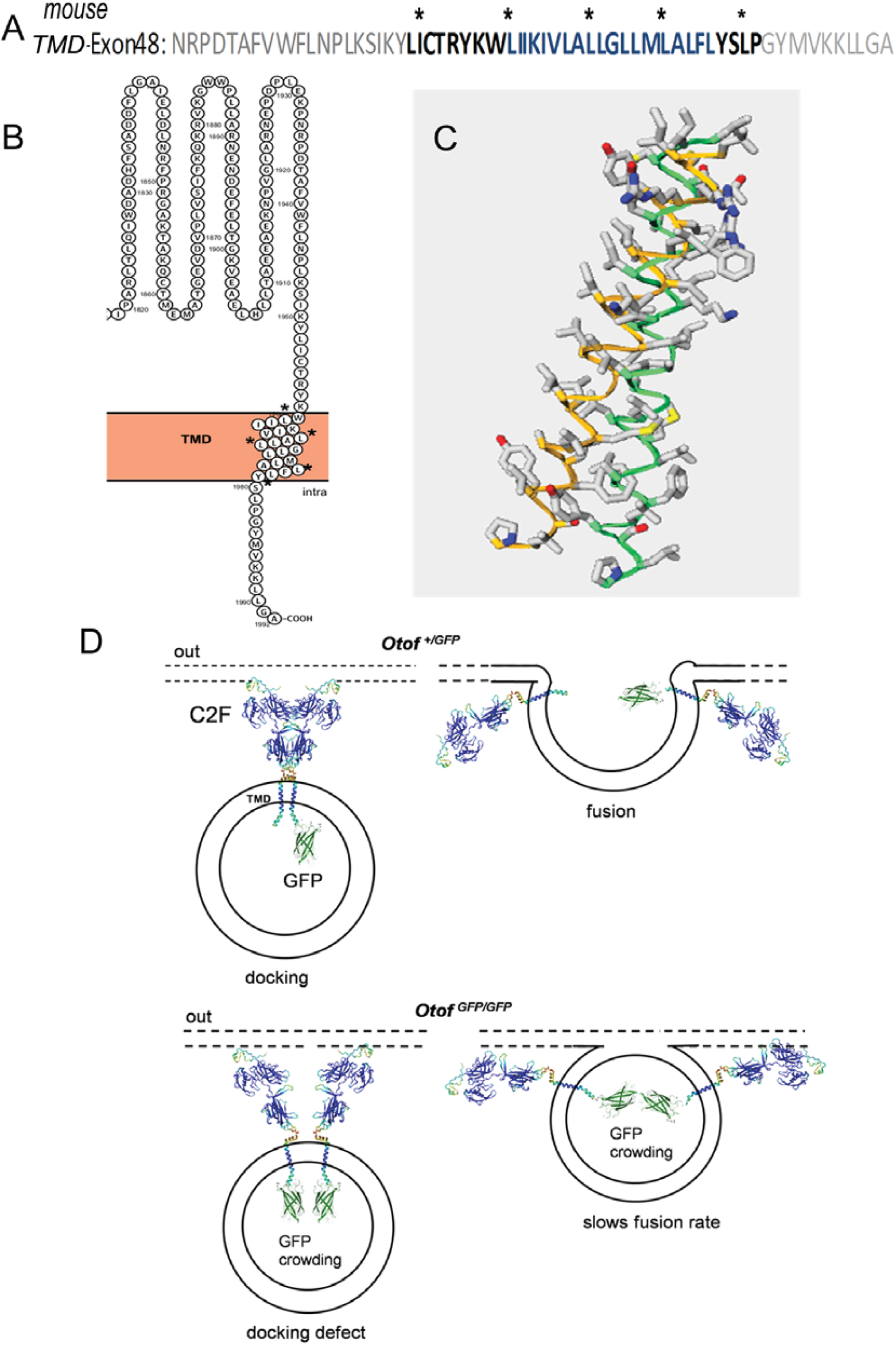
Model prediction of otoferlin dimerization through its TMD module. A) Amino acid sequence encoded by exon 48 according to Mus musculus otoferlin (Otof), transcript variant 1, mRNA (NCBI sequence identifier: NM_001100395.1). The asterisks indicate the repetitive leucine (L), isoleucine (I) heptad zipper-like structure in the juxtamembrane (black bold) and transmembrane domain (TMD, blue). B) Interactive protein feature representation using Protter analysis of the C-terminal part of otoferlin, including C2F and TMD. The asterisks indicate the L positions in the TMD. C) 3D-representation of otoferlin TMD dimer structure predicted using PREDDIMER analysis. Calculation of dimer packing probality of a-helices indicated a good F_scor_ of 3.7 (see supplemental table 1). D) Hypothetical mechanims: In *Otof ^+/GFP^* IHCs, otoferlin-TMD-GFP and otoferlin-TMD can dimerize and produce normal docking and fusion of synaptic vesicles. In *Otof ^GFP/GFP^* IHCs, because of GFP molecular crowding, otoferlin-TMD-GFP cannot properly self dimerize, then preventing or slowing-down vesicle docking and fusion.

## DISCUSSION

We generated a knock-in mouse expressing an otoferlin-GFP that enable us to track synaptic vesicles in live cochlear IHCs in both normal hearing and deaf mice. We demonstrated that the expression patterns of the otoferlin-GFP fusion protein are indistinguishable from those of native otoferlin. The heterozygote *Otof ^+/GFP^* mice are viable, healthy, and exhibit normal hearing and IHC synaptic transmission. Robust fluorescence signals could be observed in the organs of Corti of these mice, allowing for the direct detection of otoferlin-associated vesicles and for live imaging of Otof-GFP in IHCs by confocal, two-photon, or STED microscopy (Figs. 3-5). Otof-GFP was observed to accurately localize to the IHC in both *Otof^+/GFP^* and *Otof ^GFP/GFP^* mice. Moreover, neither the native otoferlin nor Otof-GFP was detected in the OHCs past the hearing onset, suggesting that the GFP tag did not disrupt the developmental regulation of otoferlin expression within the organ of Corti (Roux et al., 2006; Beurg et al., 2010). Finally, using VGLUT3 as a specific marker for IHC synaptic vesicles imaged with STED microscopy, we were able to demonstrate that Otof-GFP is targeted to both the IHC synaptic vesicle and the presynaptic plasma membrane, as previously shown in immunogold electron microscopy for native otoferlin (Roux et al., 2006; Dulon et al., 2009). Similar normal localization of synaptotagmin-GFP was found in brain synapses and synaptic vesicles (Saegusa et al., 2002; Zhang et al., 2002; Han et al., 2005).

Interestingly, heterozygote *Otof ^+/GFP^* mice exhibited normal ABRs and DPOAEs, consistent with the normal Ca^2+^-dependant exocytosis found in IHCs. FRAP experiments revealed that the mobile of GFP-associated vesicles in IHCs of *Otof ^+/GFP^* mice at room temperature is relatively low (about 20%), suggesting that a large pool of synaptic vesicles is rather immobile and had strong intracellular interactions. A similarly highly restrained population of vesicles was observed by FRAP in nerve terminals (Henkel et al., 1996; Gaffield et al., 2006). We also found that K^+^ depolarization of the IHC significantly increased the mobile fraction of Otof-GFP. This indicates that an elevation in intracellular Ca^2+^ reduces the strength of vesicle intracellular interactions within the cell, thereby ensuring a steady supply of vesicles to the active zone of the IHC ribbon synapse during sustained high rate of exocytosis, a characteristic feature of this synapse (Moser and Beutner, 2000). The coefficient of diffusion of this heightened mobile fraction remained largely unchanged during K^+^ depolarization, implying that the intracellular mobility of the synaptic vesicles is Ca^2+^ independent, consistent with previous findings from FRAP experiments in bipolar cells (Holt et al., 2004).

Surprisingly, homozygote *Otof ^GFP/GFP^* mice were profoundly deaf due to a severe impairment of IHC transmitter release, manifests in a strong reduction in the kinetics of both the RRP and SRP exocytotic components. Much like the IHCs of the constitutive *Otof* knockout mice (Tertrais et al., 2019; Roux et al., 2006), the impairment in exocytosis observed in *Otof ^GFP/GFP^* IHCs was likewise associated with an approximately 30% reduction in the number of synaptic ribbons. This confirmed the necessity of functional otoferlin for the maintenance of ribbon synapses (Stalmann et al., 2021).

Interestingly, in the absence of depolarization, FRAP experiments revealed a higher fraction mobile vesicles in the IHCs of *Otof ^GFP/GFP^* as compared to that of *Otof ^+/GFP^*, suggesting a decrease in the strength of interactions among intracellular vesicles. Importantly, previous findings in *Otof KO* IHCs, demonstrated an impact on vesicle tethering (Vogl et al., 2015; Chakrabarti et al., 2018). A comparable defect may explain the higher mobile fraction of vesicles in *Otof ^GFP/GFP^* IHCs. This could further explain the reduction in the number of synaptic vesicles connected to the active zone of the ribbon in homozygous *Otof ^GFP/GFP^* IHCs, as observed in our electron tomography analysis. Taken together, our results suggest that tagging the C-terminal otoferlin TMD with GFP impairs the protein’s function, potentially by reducing vesicular interaction and/or tethering—an indispensable step for synaptic vesicle docking and fusion. In good agreement with the study of Manchanda et al., (2021) showing that the truncation of TMD reduces membrane docking in zebrafish hair cells, our study underlines the importance of TMD.

Our study reveals the functional significance of the C-terminal portion within the intravesicular region following the otoferlin-TMD. This is in line with a clinical study showing severe hearing loss in patient bearing a substitution-mutation p.X1988RextX30, in which the stop codon in exon 48 is shifted, and 30 additional residues are added to the otoferlin TMD C-terminus intravesicular tail (Matsunaga et al. 2012). Using our in silico analysis, we have put forward a model suggesting that otoferlin function requires dimerization through its TMD (Fig.10). The available data indicates this is the most plausible assumption, since dysferlin, a close otoferlin homolog from the ferlin-1 like protein family, demonstrates TMD-mediated dimerization both in vitro and in living cells (Xu et al., 2011). Oligomerization of synaptotagmin 1, via its juxtamembrane linker, was also shown to be essential for Ca^2+^ dependent exocytosis (Bello et al., 2018; Courtney et al., 2021). It is worthwhile to note in this respect that mutations known to cause DFNB9 hearing loss lie within the juxtamembrane linker of otoferlin (p.Pro1931Leu and p.Pro1940Arg), a highly conserved domain in the ferlin family (Santarelli et al., 2021). We therefore suggest that the GFP-tag generates intravesicular molecular crowding and slows down otoferlin dimerization assembly-disassembly, a process likely essential for proper vesicle docking and fusion at the IHC ribbon active zone.

## MATERIAL AND METHODS

### Assessment of hearing function

Experiments were performed in C57BL6/J mice of either sex according to the ethical guidelines of Institut Pasteur and University of Bordeaux (French Ministry of Agriculture agreement C33-063-075; ethical committee for animal research CEEA - 050; DAP 22860-V3-2019112116537296). To record ABRs (Auditory Brainstem Responses, which represent the sound-evoked synchronous firing of the auditory cochlear nerve fibers and the activation of the subsequent central auditory relays), mice were constantly kept anesthetized under isoflurane (2.5%). The mouse was placed in a closed acoustic chamber and its body temperature was kept constant at 37°C during ABRs recording. For sound stimulus generation and data acquisition, we used a TDT RZ6/BioSigRZ system (Tucker-Davis Technologies). Click and tone-based ABR signals were averaged after the presentation of a series of 512 stimulations. Thresholds were defined as the lowest stimulus for recognizable wave-I and II. The amplitude of ABR wave-I was estimated by measuring the voltage difference between the positive (P1) and negative (N1) peak of wave-I. Sound intensities of 10–90 dB SPL in 10 dB step, were tested.

Distortion product otoacoustic emissions (DPOAEs), which originate from the electromechanical activity of the outer hair cells (OHCs), were tested by using two simultaneous continuous pure tones with frequency ratio of 1.2 (*f* _1_ = 12.73 kHz and *f* _2_ = 15.26 kHz). DPOAEs were collected with the TDT RZ6/BioSigRZ system designed to measure the level of the “cubic difference tone” 2*f*_1_–*f* _2_.

### Immunohistofluorescence

To allow rapid fixation, the round and oval windows were opened, and bone over the apical turn removed and the cochlea was perfused with 4% paraformaldehyde in PBS. After decalcification for several hours in 10% EDTA PBS, the cochlea was then post fixed in the same fixative for 30 min at 4°C. Whenever the Cav1.3 antibody was used, cochlea was fixed with 99% methanol for 20 min at-20°C. Cochlear whole-mount preparations were permeabilized with 0.3% Triton X-100 in PBS containing 20% normal Hors serum for 1 h at room temperature. The following antibodies were used: rabbit anti otoferlin (1/250), goat anti-CtBP2 (1:150; Santa Cruz Biotechnology), rabbit anti-Ca_V_1.3 (1:50; Alomone Labs), mouse anti-GluA2 (1:200; Millpore), and secondary Atto Fluor Cy5-conjugated anti-mouse, Alexa Fluor 488-conjugated anti-goat, and Atto Fluor 647-conjugated anti-rabbit IgG (1:200, sigma) Antibodies. The preparations were then washed three times in PBS, and finally mounted in one drop of Fluorsave medium (Biochem Laboratories, France). Fluorescence confocal *z*-stacks from selected cochlear regions from each ear were obtained with a Zeiss microscope using a high-resolution 63X (1.4 numerical aperture) oil-immersion objective and 4× digital zoom. Images were acquired in a 1024 × 1024 raster with 0.2 μm steps in *z*. For each genotype and for each immunofluorescence labeling and quantification, we used 72 cells from at least 6 different cochlear preparations.

### STED microscopy, Image processing and colocalization analysis

A custom-built system was used to perform dual STED microscopy (Lauterbach, 2013), using two excitation beams at 488 nm and 532 nm, and one STED (donut-shaped) beam tuned around 647 nm, all centered on the focal spot of a 100X/1.4NA objective lens (Olympus, Tokyo, Japan). STED imaging was achieved with two dyes, Atto565 and DY485XL, excited with the 532 nm and 488 nm lines, respectively (near the excitation peaks of 592 nm for Atto565 and 485 nm for DY485XL). Fluorescence was collected with an avalanche photodiode (Perkin Elmer) behind a 585/65 emission filter, encompassing the emission peaks of both dyes (around 592 nm for Atto565 and 560 nm for DY485XL). A pixel size of 20 nm and a scanning dwell time of 100 μs were used for the acquisitions.

STED images were processed by wavelet denoising and deconvolution as described (Michalski & al, 2017). The staining pattern observed in either the Atto565-or the DY485-channel images consisted in numerous resolution-limited spots representing otoferlin-GFP-and for VGLUT3-stained structures, respectively, within the IHCs. To assess the colocalization between otoferlin and VGLUT3 spots, the positions of each type of spots were first obtained by maxima detection, with a resolution typically better than the pixel size, retaining only maxima of intensity larger than 15% of the image’s maximum value. We then evaluated the distribution of the distance between a randomly chosen otoferlin spot and the closest VGLUT3 spot, which we compared with the distribution of the distance between a random VGLUT3 spot and its nearest VGLUT3 neighbor. The latter VGLUT3-VGLUT3 nearest neighbor distance distribution could be used as a proxy for the expected distribution of nearest-neighbor distances for random points of the same density as the VGLUT3 spots. A shift of the otoferlin-VGLUT3 nearest-neighbor distance distribution towards smaller distances (compared to the typical VGLUT3-VGLUT3 nearest neighbor distances) indicated colocalization of the two proteins. The significance of this shift was assessed by the Kolmogorov-Smirnov test for comparing two distributions.

### 3D-electron tomography

Cochleas were perfused with 4% PFA and 2 % glutaraldehyde in Sorensen buffer at pH 7.4, and immersed in the fixative solution for 2 h. They were then post-fixed by overnight incubation in 1% osmium tetraoxide in cacodylate buffer at 4°C, dehydrated in graded acetone concentrations, and then embedded in Spurr’s low-viscosity epoxy resin (EMS) hardened at 70°C. For tomographic analysis, thick (200 or 250 nm) sections of organs of Corti were collected on 100-mesh parallel copper grids and incubated for 30 min on each side with undiluted 15 nm gold particles conjugated with secondary anti-rabbit antibodies (UMC Utrecht, The Netherland). The grids were then stained by incubation with 4% uranyl acetate in dH_2_O for 45 min, followed by lead citrate for 3 min. The sections were transferred to an FEI *Tecnai* G2 *200kV* Transmission Electron Microscope for 3-D electron tomography. The microscope is equipped with a high resolution FEI Eagle 4K CCD camera and Digital Micrograph (Gatan) tomography acquisition software. Zero-loss tilted series (from approximately −65° to +65°, with a 1° increment). Tomographic slices were processed with a custom wavelet denoising algorithm implemented in Matlab (Mathworks), to reduce the noise while preserving fine details. The images of each tomographic tilt series were then aligned by tracking the gold particles present in the sample, and the final volume was reconstructed with a weighted back-projection algorithm, using the IMOD software (Kremer et al, 1996) for single or dual axis reconstruction.

### 3D reconstructions and pool size

Analysis of the ribbon synapses, including segmentation, 3D reconstruction, and rendering, were carried out using AMIRA software (version 5.1; Mercury Computer Systems, San Diego, CA). The contours of the ribbon, the presynaptic density of the afferent dendrite, and organelles such as mitochondria, coated pits and tubular structures, were drawn on every section. Spheres with a constant diameter were used to mark synaptic vesicles. The ribbon was defined as being the center of the presynaptic plasma membrane. For each ribbon, we counted the number of synaptic vesicles that were distant by less than 80 nm from the ribbon surface. This represented the ribbon-attached vesicle pool and considered as the slowly releasable pool (SRP). A subset of the ribbon-attached vesicles, where the distance of the vesicle was within 20 nm from the presynaptic membrane, and located below the ribbon (within 100 nm of the center of the active zone), was considered the docked pool or the readily releasable pool (RRP)—the presumed pool of vesicles that release first during a depolarization (Lenzi et al 1999, Schnee et al 2011). The value of 20 nm was chosen because t-SNARE and v-SNARE proteins have a cytosolic length of ∼ 10 nm, so SNARE interactions may occur from a distance of 20 nm from the presynaptic plasma membrane (Zenisek et al, 2000, Castorph et al., 2010). By using our ribbon reconstruction data and taking into account the SVs distribution, we estimated the total size and volume density of the ribbon-attached synaptic vesicles per ribbon, and the number and volume density of outlying cytoplasmic vesicles located within 350 nm of the ribbon’s surface. To estimate these vesicle pools in our tomographic reconstructions of ribbon synapses, we used only data sets that included more than half of the ribbon surface. Assuming that SVs attached to the ribbon are homogeneously distributed around its surface, one can estimate the vesicle pool based on the percentage of the vesicle attached to the ribbon reconstructed.

### Two-photon FRAP experiments

E*x vivo* cochlear preparation was mounted on the stage of a two-photon microscope, and superfused with an artificial perilymph solution at room temperature, or while warming the objective and chamber to 37°C. After collecting baseline images, up to four otoferlin-GFP spots were selected in different IHCs and were simultaneously bleached by a brief exposure to high-intensity laser light (∼50% of the total available intensity estimated to 2 mW at the focal plane of the objective, for an excitation wavelength of 930 nm). The bleach caused a sharp drop of fluorescence intensity, followed by a period of recovery lasting between 5 and 25 s. The FRAP image series were analyzed off-line and various parameters characterizing the fluorescence recovery were obtained: the fraction of mobile fluorescent molecules at the location of the bleached spot, the recovery half-time (*t*_1/2_), and the effective diffusion coefficient (*D*_eff_, in μm^2^/s), which was estimated by simulation of the diffusion equation after normalization of the images, according to the analysis protocol described previously for FRAP experiments on auditory hair cells (de Monvel et al., 2006). All averaged values of these parameters are given in the text as mean ± standard error.

### Tissue preparation, whole-cell patch clamp recording and capacitance measurement from IHCs

Electrophysiological recordings from IHCs were obtained in living whole-mount organs of Corti in the 20-40% normalized distance from the apex, an area coding for frequencies ranging from 8 to 16 kHz (Meyer et al., 2009). The organ of Corti (OC) was freshly dissected under binocular microscopy in an extracellular solution maintained at 4°C containing (in mM): NaCl 135; KCl 5.8; CaCl_2_ 1.3; MgCl_2_ 0.9; NaH_2_PO_4_ 0.7; Glucose 5.6; Na pyruvate 2; HEPES 10, pH 7.4, 305 mOsm. Tectorial membrane was carefully removed. The OC was placed in a recording chamber and hair cells observed with a 60x water immersion objective (CFI Fluor 60X W NIR, WD = 2.0 mm, NA = 1) attached to an upright Nikon FN1 microscope. The extracellular solution was complemented with 0.5 μM of apamin (Latoxan, Valence, France) and 0.2 μM of XE-991 (Tocris Bioscience, Lille, France) to block SK channels and KCNQ4 channels, respectively. The external Ca^2+^ concentration was increased from 1.3 mM to 5 mM in order to enhance the amplitude of Ca^2+^ currents. All experiments were carried out at room temperature (22-24 °C)

All patch clamp experiments were performed with an EPC10 amplifier controlled by pulse software Patchmaster (HEKA Elektronik, Germany). Patch pipettes were pulled with a micropipette Puller P-97 Flaming/Brown (Sutter Instrument, Novato, CA, USA) and fire polished with a Micro forge MF-830, (Narishige, Japan) to obtain a resistance range from 3 to 5 MO. Patch pipettes were filled with intracellular solution containing (in mM): CsCl 145; MgCl_2_ 1; HEPES 5; EGTA 1; TEA 20, ATP 2 and GTP 0.3, pH 7.2, 300 mOsm.

*Real time capacitance measurement*: membrane capacitance measurements (C_m_) were performed using the Lock-in amplifier Patchmaster software (HEKA) by applying a 1 kHz command sine wave (amplitude 20 mV) at holding potential (−80 mV) before and after the pulse experiment. Because capacitance measurement requires high and constant membrane resistance (R_m_), the amplitude of the sine wave was small enough to avoid activation of significant membrane current. Exocytosis (.L_Cm_) was calculated by subtracting the average capacitance measurement 60-100 ms after the depolarizing pulse (end of tail current) and the average capacitance measurement 50 ms before the pulse as baseline. Only cells with stable R_m_, leak current below 50 pA at holding potential (−80 mV) and stable series resistance below 15 MO were considered in the study.

### Caged Ca^2+^ photolysis and Ca^2+^ concentration measurement

To trigger a fast step-increase in the intracellular Ca^2+^ concentration from the caged Ca^2+^ chelator DM-nitrophen (Interchim, catalog #317210), we used a brief flash from a UV LED light source (Mic-LED 365, 128mW, Prizmatix, Givat Shmuel, Israel). The UV LED was directly connected to the epi-illumination port at the rear of the upright Nikon FN1 microscope and illumination focalized through the 60X objective (CFI Fluor 60X W NIR, WD = 2.0 mm, NA=1). Changes in [Ca^2+^]_i_ were continuously measured with a C2 confocal system and NIS-elements imaging software (Nikon, Japan) coupled to the FN1 microscope. Following patch rupture, we systematically waited for 2 min at a holding potential-of 70mV to load and equilibrate the cells with the intrapipette solution. Upon UV photolysis, [Ca^2+^]_i_, continuously measured with a C2 confocal system and NIS-Elements imaging software (Nikon) coupled to the FN1 microscope, reached a peak of 20 ± 5 µM within 15–20 ms (Vincent et al., 2014). For Ca^2+^ uncaging, in which membrane capacitance was continuously recorded, patch pipettes were filled with the following solution (in mM): CsCl 145; HEPES 5; TEA 20; DM-nitrophen 10; CaCl_2_ 10.

### Statistical Analysis

Results were analyzed with OriginPro 9.1software (OriginLab, Northampton, USA), and in Matlab (The Mathworks, Natik, USA). Data were tested for normal distribution using the Shapiro–Wilk normality test, and parametric or nonparametric tests were applied accordingly. For statistical analyses with two data sets, two-tailed unpaired *t*-tests or two-sided Mann– Whitney tests were used. For comparisons of more than two data sets, one-way ANOVA or two-way ANOVA followed by Tukey multiple comparison test were used when data was normally distributed. If a data set was not normally distributed Kruskal–Wallis tests followed by Dunn multiple comparison tests, respectively, was used. When more than two data sets were compared and significant differences were found, reported *p*-values correspond to the *post-hoc* multiple comparison test. Values of *p* < 0.05 were considered significant. Results are expressed as mean ± SEM or SD as indicated in the text.

## Acknowledgments

We would like to thank Anna Sartori-Rupp of the “Plate-Forme de Microscopie ultrastructurale of the Pasteur Institute. Part of the confocal fluorescence microscopy was done in the Bordeaux Imaging Center, a service unit of the CNRS-INSERM and Bordeaux University, member of the national infrastructure France BioImaging. We are also grateful for their help in some experiments of the study to Sylvie Nouaille, Philippe Vincent, Margot Tertrais and Thibault Peineau. This work was supported by the French National Research Agency which is funding the future investment program entitled RHU AUDINNOVE, ANR-18-RHUS-0007 to SS”, programme EARGENCURE (ANR-17-CE18-0027 to SS). This work was in large part financed by the Fondation pour l’Audition (FPA grant number: n° FPA IDA09 to DD and FPA IDA08 to SS).

## SUPPLEMENTARY

**Figure S1:**
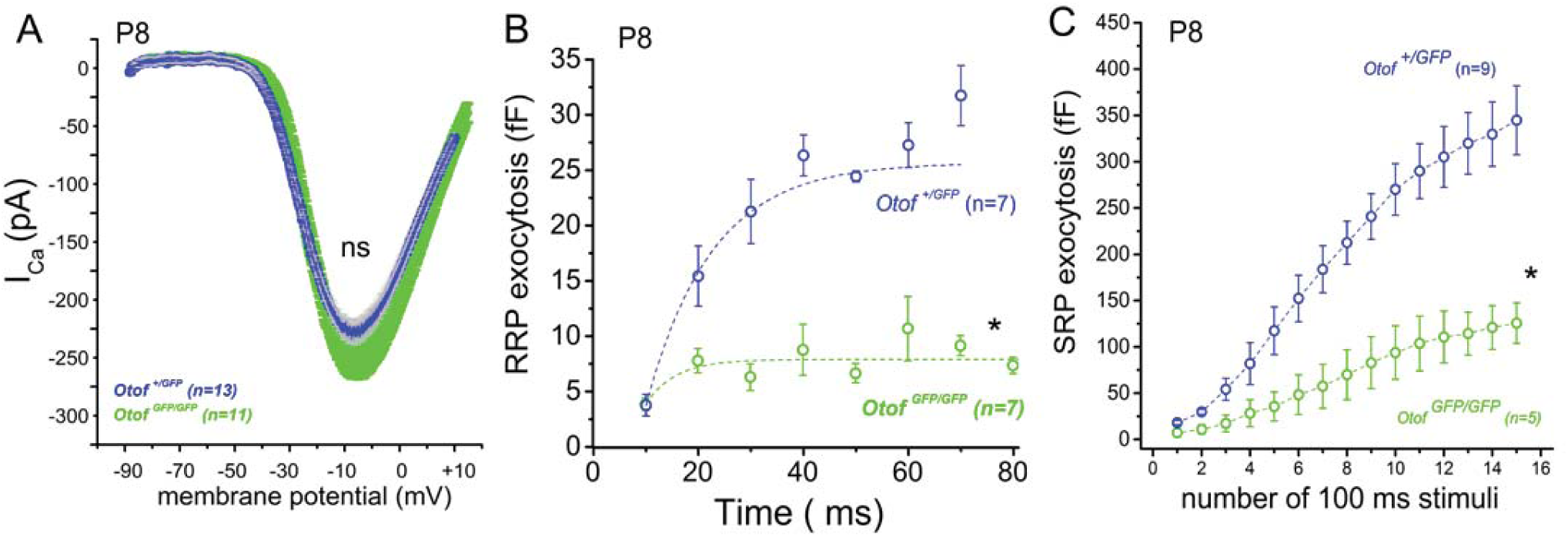
Exocytosis defect in IHCs from immature prehearing P8 *Otof ^GFFP/GFP^* mice. A) Comparable Ca^2+^ currents evoked by a voltage-ramp protocol in *Otof ^+/GFP^* (blue) and *Otof^GFP/GFP^* (green). B-C) Kinetics of RRP and SRP *Otof^GFP/GFP^* in IHCs, using similar protocols described in figures 6 and 7, were largely decreased as compared to *Otof ^+/GFP^* IHCs (blue). Values are mean with SEM and asterisks indicate significant difference with p<0.05 while ns indicates not significantly different (two-way ANOVA).

**Supp table 1:** Prediction results using PREDDIMER of transmembrane a-helical dimer conformations of the mouse otoferlin TMD. *F_Sco_*_r=_ *Pack*.(*Int + Env*) indicate the dimer packing quality. *Pack* is the term corresponding to a relative number of atoms packed within the structures, *Int* is the term accounting complementarity of hydrophobic properties on the helix—helix interface, *Env* is the term estimating correspondence of polar and structural properties of the dimer surface exposed to lipid environment.

| Rank | $F_{\text{score}}$ | Crossing angle $\chi$, deg | Rotation angle $\alpha_1$, deg | Rotation angle $\alpha_2$, deg |
| --- | --- | --- | --- | --- |
| 1 | 3.704 | -15.8 | 338 | 458 |
| 2 | 3.19 | 60 | 488 | 488 |
| 3 | 2.906 | -11.5 | 452 | 314 |
| 4 | 2.305 | 30.1 | 404 | 392 |
| 5 | 2.244 | -40.1 | -100 | -100 |
| 6 | 2.185 | -55 | -124 | -130 |
| 7 | 2.121 | -30.3 | 194 | 194 |
| 8 | 1.21 | 25 | 170 | 170 |

## REFERENCES

Bello OD, Jouannot O, Chaudhuri A, Stroeva E, Coleman J, Volynski KE, Rothman JE, Krishnakumar SS. Synaptotagmin oligomerization is essential for calcium control of regulated exocytosis. Proc Natl Acad Sci U S A. 2018 Aug 7;115(32):E7624–E7631. doi: 10.1073/pnas.1808792115. Epub 2018 Jul 23. PMID: 30038018; PMCID: PMC6094142. Format:

Brandt A, Khimich D, Moser T. Few CaV1.3 channels regulate the exocytosis of a synaptic vesicle at the hair cell ribbon synapse. J Neurosci. 2005 Dec 14;25(50):11577–85. doi: 10.1523/JNEUROSCI.3411-05.2005. PMID: 16354915; PMCID: PMC6726013.

Beurg M, Safieddine S, Roux I, Bouleau Y, Petit C, Dulon D. Calcium-and otoferlin-dependent exocytosis by immature outer hair cells. J Neurosci. 2008 Feb 20;28(8):1798–803. doi: 10.1523/JNEUROSCI.4653-07.2008. PMID: 18287496; PMCID: PMC6671446.

Calvet C, Peineau T, Benamer N, Cornille M, Lelli A, Plion B, Lahlou G, Fanchette J, Nouaille S, Boutet de Monvel J, Estivalet A, Jean P, Michel V, Sachse M, Michalski N, Avan P, Petit C, Dulon D, Safieddine S. The SNARE protein SNAP-25 is required for normal exocytosis at auditory hair cell ribbon synapses. iScience. 2022 Nov 22;25(12):105628.

Castorph S, Riedel D, Arleth L, Sztucki M, Jahn R, Holt M, Salditt T. Structure parameters of synaptic vesicles quantified by small-angle x-ray scattering. Biophys J. 2010 Apr 7;98(7):1200–8. doi: 10.1016/j.bpj.2009.12.4278. PMID: 20371319; PMCID: PMC2849067.

Chakrabarti R, Michanski S, Wichmann C. Vesicle sub-pool organization at inner hair cell ribbon synapses. EMBO Rep. 2018 Nov;19(11):e44937. doi: 10.15252/embr.201744937. Epub 2018 Sep 10. PMID: 30201800; PMCID: PMC6216280.

Courtney KC, Vevea JD, Li Y, Wu Z, Zhang Z, Chapman ER. Synaptotagmin 1 oligomerization via the juxtamembrane linker regulates spontaneous and evoked neurotransmitter release. Proc Natl Acad Sci U S A. 2021 Nov 30;118(48):e2113859118. doi: 10.1073/pnas.2113859118. PMID: 34810248; PMCID: PMC8694047.

Cymer,F. and Schneider,D. (2010) Transmembrane helix-helix interactions involved in ErbB receptor signaling. Cell Adh. Migr., 4, 299–312.

de Caprona MD, Beisel KW, Nichols DH, Fritzsch B. Partial behavioral compensation is revealed in balance tasked mutant mice lacking otoconia. Brain Res Bull. 2004 Dec 15;64(4):289–301. doi: 10.1016/j.brainresbull.2004.08.004. PMID: 15561463.

de Monvel JB, Brownell WE, Ulfendahl M. Lateral diffusion anisotropy and membrane lipid/skeleton interaction in outer hair cells. Biophys J. 2006 Jul 1;91(1):364–81. doi: 10.1529/biophysj.105.076331. Epub 2006 Apr 7. Erratum in: Biophys J. 2018 Jun 5;114(11):2756. PMID: 16603502; PMCID: PMC1479061.

Dulon D, Safieddine S, Jones SM, Petit C. Otoferlin is critical for a highly sensitive and linear calcium-dependent exocytosis at vestibular hair cell ribbon synapses. J Neurosci. 2009 Aug 26;29(34):10474–87. doi: 10.1523/JNEUROSCI.1009-09.2009. PMID: 19710301; PMCID: PMC2966717.

Gaffield MA, Rizzoli SO, Betz WJ. Mobility of synaptic vesicles in different pools in resting and stimulated frog motor nerve terminals. Neuron. 2006 Aug 3;51(3):317–25. doi: 10.1016/j.neuron.2006.06.031. PMID: 16880126.

Glowatzki E, Fuchs PA. Transmitter release at the hair cell ribbon synapse. Nat Neurosci. 2002 Feb;5(2):147–54. doi: 10.1038/nn796. PMID: 11802170.

Goutman JD, Glowatzki E. Time course and calcium dependence of transmitter release at a single ribbon synapse. Proc Natl Acad Sci U S A. 2007 Oct 9;104(41):16341–6. doi: 10.1073/pnas.0705756104. Epub 2007 Oct 2. PMID: 17911259; PMCID: PMC2042208.

Gurezka R, Rico Laage, Bettina Brosig, Dieter Langosch,A Heptad Motif of Leucine Residues Found in Membrane Proteins Can Drive Self-assembly of Artificial Transmembrane Segments*, Journal of Biological Chemistry, Volume 274, Issue 14, 1999, Pages 9265-9270, ISSN 0021-9258, 10.1074/jbc.274.14.9265

Han W, Rhee JS, Maximov A, Lin W, Hammer RE, Rosenmund C, Südhof TC. C-terminal ECFP fusion impairs synaptotagmin 1 function: crowding out synaptotagmin 1. J Biol Chem. 2005 Feb 11;280(6):5089–100. doi: 10.1074/jbc.M408757200. Epub 2004 Nov 23. PMID: 15561725.

Hell SW, Wichmann J. Breaking the diffraction resolution limit by stimulated emission: stimulated-emission-depletion fluorescence microscopy. Opt Lett. 1994 Jun 1;19(11):780–2. doi: 10.1364/ol.19.000780. PMID: 19844443.

Henkel AW, Simpson LL, Ridge RM, Betz WJ. Synaptic vesicle movements monitored by fluorescence recovery after photobleaching in nerve terminals stained with FM1-43. J Neurosci. 1996 Jun 15;16(12):3960–7. doi: 10.1523/JNEUROSCI.16-12-03960.1996. PMID: 8656290; PMCID: PMC6578606.

Heidrych P, Zimmermann U, Bress A, Pusch CM, Ruth P, Pfister M, Knipper M, Blin N. Rab8b GTPase, a protein transport regulator, is an interacting partner of otoferlin, defective in a human autosomal recessive deafness form. Hum Mol Genet. 2008 Dec 1;17(23):3814–21. doi: 10.1093/hmg/ddn279. Epub 2008 Sep 4. PMID: 18772196.

Heidrych P, Zimmermann U, Kuhn S, Franz C, Engel J, Duncker SV, Hirt B, Pusch CM, Ruth P, Pfister M, Marcotti W, Blin N, Knipper M. Otoferlin interacts with myosin VI: implications for maintenance of the basolateral synaptic structure of the inner hair cell. Hum Mol Genet. 2009 Aug 1;18(15):2779–90. doi: 10.1093/hmg/ddp213. Epub 2009 May 5. PMID: 19417007.

Holt M, Cooke A, Neef A, Lagnado L. High mobility of vesicles supports continuous exocytosis at a ribbon synapse. Curr Biol. 2004 Feb 3;14(3):173–83. doi: 10.1016/j.cub.2003.12.053. PMID: 14761649.

Iwasa YI, Nishio SY, Sugaya A, Kataoka Y, Kanda Y, Taniguchi M, Nagai K, Naito Y, Ikezono T, Horie R, Sakurai Y, Matsuoka R, Takeda H, Abe S, Kihara C, Ishino T, Morita SY, Iwasaki S, Takahashi M, Ito T, Arai Y, Usami SI. OTOF mutation analysis with massively parallel DNA sequencing in 2,265 Japanese sensorineural hearing loss patients. PLoS One. 2019 May 16;14(5):e0215932. doi: 10.1371/journal.pone.0215932. PMID: 31095577; PMCID: PMC6522017.

Jumper J, Evans R, Pritzel A, Green T, Figurnov M, Ronneberger O, Tunyasuvunakool K, Bates R, Žídek A, Potapenko A, Bridgland A, Meyer C, Kohl SAA, Ballard AJ, Cowie A, Romera-Paredes B, Nikolov S, Jain R, Adler J, Back T, Petersen S, Reiman D, Clancy E, Zielinski M, Steinegger M, Pacholska M, Berghammer T, Bodenstein S, Silver D, Vinyals O, Senior AW, Kavukcuoglu K, Kohli P, Hassabis D. Highly accurate protein structure prediction with AlphaFold. Nature. 2021 Aug;596(7873):583-589. doi: 10.1038/s41586-021-03819-2. Epub 2021 Jul 15. PMID: 34265844; PMCID: PMC8371605.

Landschulz WH, Johnson PF, McKnight SL. The leucine zipper: a hypothetical structure common to a new class of DNA binding proteins. Science. 1988 Jun 24;240(4860):1759-64. doi: 10.1126/science.3289117. PMID: 3289117.

Lauterbach MA, Guillon M, Soltani A, Emiliani V. STED microscope with spiral phase contrast. Sci Rep. 2013;3:2050. doi: 10.1038/srep02050. PMID: 23787399; PMCID: PMC3689173.

Leclère JC, Dulon D. Otoferlin as a multirole Ca^2+^ signaling protein: from inner ear synapses to cancer pathways. Front Cell Neurosci. 2023 Jul 19;17:1197611.

Lenzi D and von Gersdorff H Structure suggests function the case for synaptic ribbons as exocytotic nanomachines. Bioessays. 2001; 23: 831–840.

Lenzi D, Crum J, Ellisman MH, Roberts WM. Depolarization redistributes synaptic membrane and creates a gradient of vesicles on the synaptic body at a ribbon synapse. Neuron. 2002 Nov 14;36(4):649–59. doi: 10.1016/s0896-6273(02)01025-5. PMID: 12441054.

Li Q, Wong YL, Huang Q, Kang C. Structural insight into the transmembrane domain and the juxtamembrane region of the erythropoietin receptor in micelles. Biophys J. 2014 Nov 18;107(10):2325–36. doi: 10.1016/j.bpj.2014.10.013. PMID: 25418301; PMCID: PMC4241451.

Manchanda A, Bonventre JA, Bugel SM, Chatterjee P, Tanguay R, Johnson CP. Truncation of the otoferlin transmembrane domain alters the development of hair cells and reduces membrane docking. Mol Biol Cell. 2021 Jul 1;32(14):1293–1305. doi: 10.1091/mbc.E20-10-0657. Epub 2021 May 12. PMID: 33979209; PMCID: PMC8351550.

Matsunaga T, Mutai H, Kunishima S, Namba K, Morimoto N, Shinjo Y, Arimoto Y, Kataoka Y, Shintani T, Morita N, Sugiuchi T, Masuda S, Nakano A, Taiji H, Kaga K. A prevalent founder mutation and genotype-phenotype correlations of OTOF in Japanese patients with auditory neuropathy. Clin Genet. 2012 Nov;82(5):425–32. doi: 10.1111/j.1399-0004.2012.01897.x. Epub 2012 Jun 1. PMID: 22575033.

Michalski N, Goutman JD, Auclair SM, Boutet de Monvel J, Tertrais M, Emptoz A, Parrin A, Nouaille S, Guillon M, Sachse M, Ciric D, Bahloul A, Hardelin JP, Sutton RB, Avan P, Krishnakumar SS, Rothman JE, Dulon D, Safieddine S, Petit C. Otoferlin acts as a Ca^2+^ sensor for vesicle fusion and vesicle pool replenishment at auditory hair cell ribbon synapses. Elife. 2017 Nov 7;6:e31013. doi: 10.7554/eLife.31013. PMID: 29111973; PMCID: PMC5700815.

Moser T, Beutner D. Kinetics of exocytosis and endocytosis at the cochlear inner hair cell afferent synapse of the mouse. Proc Natl Acad Sci U S A. 2000 Jan 18;97(2):883–8. doi: 10.1073/pnas.97.2.883. PMID: 10639174; PMCID: PMC15425.

Obholzer N, Wolfson S, Trapani JG, Mo W, Nechiporuk A, Busch-Nentwich E, Seiler C, Sidi S, Söllner C, Duncan RN, Boehland A, Nicolson T. Vesicular glutamate transporter 3 is required for synaptic transmission in zebrafish hair cells. J Neurosci. 2008 Feb 27;28(9):2110–8. doi: 10.1523/JNEUROSCI.5230-07.2008. PMID: 18305245; PMCID: PMC6671858.

Omasits U, Ahrens CH, Müller S, Wollscheid B Protter: interactive protein feature visualization and integration with experimental proteomic data.. Bioinformatics. 2014 Mar 15;30(6):884–6. doi: 10.1093/bioinformatics/btt607.

Pangrsic T, Lasarow L, Reuter K, Takago H, Schwander M, Riedel D, Frank T, Tarantino LM, Bailey JS, Strenzke N, Brose N, Müller U, Reisinger E, Moser T. Hearing requires otoferlin-dependent efficient replenishment of synaptic vesicles in hair cells. Nat Neurosci. 2010 Jul;13(7):869–76. doi: 10.1038/nn.2578. Epub 2010 Jun 20. PMID: 20562868.

Polyansky A.A., Volynsky P.E., Efremov R.G. (2012). Multistate organization of transmembrane helical protein dimers governed by the host membrane. J. Am. Chem. Soc. 134(35), 14390–14400

Polyansky AA, Chugunov AO, Volynsky PE, Krylov NA, Nolde DE, Efremov RG. PREDDIMER: a web server for prediction of transmembrane helical dimers. Bioinformatics. 2014 Mar 15;30(6):889–90. doi: 10.1093/bioinformatics/btt645. Epub 2013 Nov 7. PMID: 24202542.

Roux I, Safieddine S, Nouvian R, Grati M, Simmler MC, Bahloul A, Perfettini I, Le Gall M, Rostaing P, Hamard G, Triller A, Avan P, Moser T, Petit C. Otoferlin, defective in a human deafness form, is essential for exocytosis at the auditory ribbon synapse. Cell. 2006 Oct 20;127(2):277–89. doi: 10.1016/j.cell.2006.08.040. PMID: 17055430.

Ruel J, Emery S, Nouvian R, Bersot T, Amilhon B, Van Rybroek JM, Rebillard G, Lenoir M, Eybalin M, Delprat B, Sivakumaran TA, Giros B, El Mestikawy S, Moser T, Smith RJ, Lesperance MM, Puel JL. Impairment of SLC17A8 encoding vesicular glutamate transporter-3, VGLUT3, underlies nonsyndromic deafness DFNA25 and inner hair cell dysfunction in null mice. Am J Hum Genet. 2008 Aug;83(2):278–92. doi: 10.1016/j.ajhg.2008.07.008. PMID: 18674745; PMCID: PMC2495073.

Rutherford MA, Bhattacharyya A, Xiao M, Cai HM, Pal I, Rubio ME. GluA3 subunits are required for appropriate assembly of AMPAR GluA2 and GluA4 subunits on cochlear afferent synapses and for presynaptic ribbon modiolar-pillar morphology. Elife. 2023 Jan 17;12:e80950. doi: 10.7554/eLife.80950. PMID: 36648432; PMCID: PMC9891727.

Saegusa C, Fukuda M, Mikoshiba K. Synaptotagmin V is targeted to dense-core vesicles that undergo calcium-dependent exocytosis in PC12 cells. J Biol Chem. 2002 Jul 5;277(27):24499–505. doi: 10.1074/jbc.M202767200. Epub 2002 May 2. PMID: 12006594.

Safieddine S, El-Amraoui A, Petit C. The auditory hair cell ribbon synapse: from assembly to function. Annu Rev Neurosci. 2012;35:509–28. doi: 10.1146/annurev-neuro-061010-113705. PMID: 22715884.

Santarelli R, Scimemi P, Costantini M, Domínguez-Ruiz M, Rodríguez-Ballesteros M, Del Castillo I. Cochlear Synaptopathy due to Mutations in OTOF Gene May Result in Stable Mild Hearing Loss and Severe Impairment of Speech Perception. Ear Hear. 2021 Nov-Dec 01;42(6):1627-1639. doi: 10.1097/AUD.0000000000001052. PMID: 33908410; PMCID: PMC9973442.

Seal RP, Akil O, Yi E, Weber CM, Grant L, Yoo J, Clause A, Kandler K, Noebels JL, Glowatzki E, Lustig LR, Edwards RH. Sensorineural deafness and seizures in mice lacking vesicular glutamate transporter 3. Neuron. 2008 Jan 24;57(2):263–75. doi: 10.1016/j.neuron.2007.11.032. PMID: 18215623; PMCID: PMC2293283.

Sergeyenko Y, Lall K, Liberman MC, Kujawa SG. Age-related cochlear synaptopathy: an early-onset contributor to auditory functional decline. J Neurosci. 2013 Aug 21;33(34):13686–94. doi: 10.1523/JNEUROSCI.1783-13.2013. PMID: 23966690; PMCID: PMC3755715.

Schmitz F, Königstorfer A, Südhof TC. RIBEYE, a component of synaptic ribbons: a protein’s journey through evolution provides insight into synaptic ribbon function. Neuron. 2000 Dec;28(3):857–72. doi: 10.1016/s0896-6273(00)00159-8. PMID: 11163272.

Schug N, Braig C, Zimmermann U, Engel J, Winter H, Ruth P, Blin N, Pfister M, Kalbacher H, Knipper M. Differential expression of otoferlin in brain, vestibular system, immature and mature cochlea of the rat. Eur J Neurosci. 2006 Dec;24(12):3372–80. doi: 10.1111/j.1460-9568.2006.05225.x. PMID: 17229086.

Smith CA, Sjostrand FS. Structure of the nerve endings on the external hair cells of the guinea pig cochlea as studied by serial sections. J Ultrastruct Res. 1961 Dec;5:523–56. doi: 10.1016/s0022-5320(61)80025-7. PMID: 13914158.

Stalmann U, Franke AJ, Al-Moyed H, Strenzke N, Reisinger E. Otoferlin Is Required for Proper Synapse Maturation and for Maintenance of Inner and Outer Hair Cells in Mouse Models for DFNB9. Front Cell Neurosci. 2021 Jul 14;15:677543. doi: 10.3389/fncel.2021.677543. PMID: 34335185; PMCID: PMC8316924.

Tertrais M, Bouleau Y, Emptoz A, Belleudy S, Sutton RB, Petit C, Safieddine S, Dulon D. Viral Transfer of Mini-Otoferlins Partially Restores the Fast Component of Exocytosis and Uncovers Ultrafast Endocytosis in Auditory Hair Cells of Otoferlin Knock-Out Mice. J Neurosci. 2019 May 1;39(18):3394–3411. doi: 10.1523/JNEUROSCI.1550-18.2018. Epub 2019 Mar 4. PMID: 30833506; PMCID: PMC6495124.

Uthaiah RC, Hudspeth AJ. Molecular anatomy of the hair cell’s ribbon synapse. J Neurosci. 2010 Sep 15;30(37):12387–99. doi: 10.1523/JNEUROSCI.1014-10.2010. PMID: 20844134; PMCID: PMC2945476.

Varadi M, Anyango S, Deshpande M, Nair S, Natassia C, Yordanova G, Yuan D, Stroe O, Wood G, Laydon A, Žídek A, Green T, Tunyasuvunakool K, Petersen S, Jumper J, Clancy E, Green R, Vora A, Lutfi M, Figurnov M, Cowie A, Hobbs N, Kohli P, Kleywegt G, Birney E, Hassabis D, Velankar S. AlphaFold Protein Structure Database: massively expanding the structural coverage of protein-sequence space with high-accuracy models. Nucleic Acids Res. 2022 Jan 7;50(D1):D439–D444. doi: 10.1093/nar/gkab1061. PMID: 34791371; PMCID: PMC8728224.

Vincent PF, Bouleau Y, Safieddine S, Petit C, Dulon D. Exocytotic machineries of vestibular type I and cochlear ribbon synapses display similar intrinsic otoferlin-dependent Ca2+ sensitivity but a different coupling to Ca2+ channels. J Neurosci. 2014 Aug 13;34(33):10853–69. doi: 10.1523/JNEUROSCI.0947-14.2014. PMID: 25122888; PMCID: PMC6705247.

Vincent PF, Bouleau Y, Charpentier G, Emptoz A, Safieddine S, Petit C, Dulon D. Different Ca_V_1.3 Channel Isoforms Control Distinct Components of the Synaptic Vesicle Cycle in Auditory Inner Hair Cells. J Neurosci. 2017 Mar 15;37(11):2960–2975. doi: 10.1523/JNEUROSCI.2374-16.2017. Epub 2017 Feb 13. PMID: 28193694; PMCID: PMC6596729.

Vincent PFY, Cho S, Tertrais M, Bouleau Y, von Gersdorff H, Dulon D. Clustered Ca^2+^ Channels Are Blocked by Synaptic Vesicle Proton Release at Mammalian Auditory Ribbon Synapses. Cell Rep. 2018 Dec 18;25(12):3451–3464.e3. doi: 10.1016/j.celrep.2018.11.072. PMID: 30566869; PMCID: PMC6365105.

Vogl C, Cooper BH, Neef J, Wojcik SM, Reim K, Reisinger E, Brose N, Rhee JS, Moser T, Wichmann C. Unconventional molecular regulation of synaptic vesicle replenishment in cochlear inner hair cells. J Cell Sci. 2015 Feb 15;128(4):638–44. doi: 10.1242/jcs.162099. Epub 2015 Jan 20. PMID: 25609709.

Vogl C, Panou I, Yamanbaeva G, Wichmann C, Mangosing SJ, Vilardi F, Indzhykulian AA, Pangrsic T, Santarelli R, Rodriguez-Ballesteros M, Weber T, Jung S, Cardenas E, Wu X, Wojcik SM, Kwan KY, Del Castillo I, Schwappach B, Strenzke N, Corey DP, Lin SY, Moser T. Tryptophan-rich basic protein (WRB) mediates insertion of the tail-anchored protein otoferlin and is required for hair cell exocytosis and hearing. EMBO J. 2016 Dec 1;35(23):2536–2552. doi: 10.15252/embj.201593565. Epub 2016 Jul 25. PMID: 27458190; PMCID: PMC5283584.

Vona B, Rad A, Reisinger E. The Many Faces of DFNB9: Relating *OTOF* Variants to Hearing Impairment. Genes (Basel). 2020 Nov 26;11(12):1411. doi: 10.3390/genes11121411. PMID: 33256196; PMCID: PMC7768390.

Westphal V, Rizzoli SO, Lauterbach MA, Kamin D, Jahn R, Hell SW. Video-rate far-field optical nanoscopy dissects synaptic vesicle movement. Science. 2008 Apr 11;320(5873):246-9. doi: 10.1126/science.1154228. Epub 2008 Feb 21. PMID: 18292304.

Willig KI, Kellner RR, Medda R, Hein B, Jakobs S, Hell SW. Nanoscale resolution in GFP-based microscopy. Nat Methods. 2006 Sep;3(9):721–3. doi: 10.1038/nmeth922. PMID: 16896340.

Xu L, Pallikkuth S, Hou Z, Mignery GA, Robia SL, Han R. Dysferlin forms a dimer mediated by the C2 domains and the transmembrane domain in vitro and in living cells. PLoS One. 2011;6(11):e27884. doi: 10.1371/journal.pone.0027884. Epub 2011 Nov 14. PMID: 22110769; PMCID: PMC3215728.

Yasunaga S, Grati M, Cohen-Salmon M, El-Amraoui A, Mustapha M, Salem N, El-Zir E, Loiselet J, Petit C. A mutation in OTOF, encoding otoferlin, a FER-1-like protein, causes DFNB9, a nonsyndromic form of deafness. Nat Genet. 1999 Apr;21(4):363–9. doi: 10.1038/7693. PMID: 10192385.

Zenisek D, Steyer JA, Almers W. Transport, capture and exocytosis of single synaptic vesicles at active zones. Nature. 2000 Aug 24;406(6798):849-54. doi: 10.1038/35022500. PMID: 10972279.

Zhang YQ, Rodesch CK, Broadie K. Living synaptic vesicle marker: synaptotagmin-GFP. Genesis. 2002 Sep-Oct;34(1-2):142-5. doi: 10.1002/gene.10144. PMID: 12324970.

